# evSeq: Cost-Effective Amplicon Sequencing of Every Variant in a Protein Library

**DOI:** 10.1101/2021.11.18.469179

**Authors:** Bruce J. Wittmann, Kadina E. Johnston, Patrick J. Almhjell, Frances H. Arnold

## Abstract

Widespread availability of protein sequence-fitness data would revolutionize both our biochemical understanding of proteins and our ability to engineer them. Unfortunately, even though thousands of protein variants are generated and evaluated for fitness during a typical protein engineering campaign, most are never sequenced, leaving a wealth of potential sequence-fitness information untapped. This largely stems from the fact that sequencing is unnecessary for many protein engineering strategies; the added cost and effort of sequencing is thus unjustified. Here, we present every variant sequencing (evSeq), an efficient protocol for sequencing a variable region within every variant gene produced during a protein engineering campaign at a cost of cents per variant. Execution of evSeq is simple, requires no sequencing experience to perform, relies only on resources and services typically available to biology labs, and slots neatly into existing protein engineering workflows. Analysis of evSeq data is likewise made simple by its accompanying software (found at github.com/fhalab/evSeq, documentation at fhalab.github.io/evSeq), which can be run on a personal laptop and was designed to be accessible to users with no computational experience. Low-cost and easy to use, evSeq makes collection of extensive protein variant sequence-fitness data practical.

## INTRODUCTION

Engineered proteins are valuable tools across the biological and chemical sciences and have revolutionized industries ranging from food to fuels, pharmaceuticals, and textiles by providing green and efficient protein solutions to challenging chemical problems.^1^ Over the course of a protein engineering campaign, hundreds to thousands or more protein variants will be constructed and have their fitnesses (level of, e.g., thermostability, catalytic activity, substrate binding, etc.) evaluated. Notably, sequence information is typically not gathered alongside the functional information, even though it could provide useful biochemical insight.^2–4^ This is largely because many engineering strategies can be applied without sequencing. For example, during a typical directed evolution (DE) experiment, often only the best-performing variant or variants are sequenced in each round of mutagenesis and screening; sequencing every variant is viewed as an unnecessary expense. Given the massive amount of functional data gathered during a typical DE campaign, however, if sequencing *were* performed for the variants generated during these experiments, the resultant large datasets of sequence-fitness information could be revolutionary for biological, biochemical, and biocatalytic research. This is especially true for data-driven protein engineering strategies such as machine learning (ML), the development of which has benefitted tremendously from large sequence-fitness datasets made available by strategies like deep mutational scanning (DMS) and in databases like ProtaBank.^5–16^

Unfortunately, the standard sequencing strategy employed during DE—Sanger sequencing—is too expensive for sequencing all variants tested during a round of evolution.^17^ Sanger sequencing is ubiquitous due to ease of sample preparation, accessibility of sequencing providers, and low cost. However, this cost scales linearly with the number of samples (Supplemental Figure S1). Thus, while the cost of sequencing just the top variants in a round of DE is minor, sequencing the hundreds or thousands of variants generated over the full engineering endeavor is not. Ideally, any new method for sequencing every variant produced during a protein engineering campaign would be comparable in cost and effort to that of sequencing just the top variants by Sanger sequencing. Here we present a method that accomplishes this goal. The method, which we call every variant sequencing (evSeq), expands on services made available by multiplexed next-generation sequencing (NGS) providers to allow amplicon sequencing of a region of interest within every variant produced during a round of DE at a cost of cents per variant.^18,19^ Sample preparation for evSeq is simple, and the method requires no experience with NGS to perform, relies only on resources and services typically available to biology labs, and slots neatly into existing protein engineering experimental workflows. The accompanying software for analysis of evSeq data (found at github.com/fhalab/evSeq, documentation at fhalab.github.io/evSeq) was designed to be accessible to users with no computational experience and can be run on a personal laptop.

In this paper, we detail the development, protocol, and potential applications of evSeq. We begin by describing the strategies employed by evSeq to extend multiplexed NGS for sequencing protein variant libraries in a way that reduces both cost and effort. We then describe the wet-lab protocol of evSeq sample preparation, focusing on how it can be completed without disrupting an existing protein engineering workflow. Next, we discuss the features of the evSeq software before finally presenting two case studies that highlight potential applications of evSeq. In particular, we highlight how (1) the sequence-fitness data from evSeq can provide valuable information about the quality of variant libraries and the functional screen as well as how mutations modulate protein activity, and how (2) the data generated from evSeq can be used to implement ML for protein engineering. We designed evSeq for use as a routine procedure in many protein/enzyme assays (especially DE and protein engineering experiments leveraging mutagenesis strategies that target specific sites or a segment of the sequence). We believe that widespread adoption of evSeq—and the resultant datasets generated— will be invaluable for future ML-guided protein engineering and will help us better understand protein sequence-fitness relationships.

## RESULTS AND DISCUSSION

### evSeq expands on commercially available multiplexed next-generation sequencing

Unlike Sanger sequencing, which outputs a single chromatogram that represents the population of DNA in a sequenced sample, NGS outputs millions of individual DNA reads that represent a random draw from the population of DNA in the sequenced sample.^18^ Confidence in NGS sequencing results is largely determined by the sequencing “coverage”, which for the purposes of this paper is defined as the number of returned reads that map to a specific nucleotide on a reference sequence. Higher coverage enables more confident identification of mutations relative to a reference sequence as the increased redundancy allows distinguishing between true sequence mutations and errors that arise during library preparation, clustering, or sequencing.

A single NGS run is roughly three orders of magnitude more expensive than a Sanger sequencing run, but because the run outputs millions of reads this cost can be spread over multiple samples using a technique known as “multiplexed NGS” (Supplemental Figure S1). In multiplexed NGS, each submitted sample is tagged with a “molecular barcode”—a unique piece of DNA that encodes the sample’s original identity—before all samples are sequenced together in the same NGS run.^19–25^ Post sequencing, the barcodes are used to assign individual reads to individual samples. Multiplexed NGS can be outsourced just like Sanger sequencing, and sequencing providers typically charge tens of dollars per sample in a multiplexed sequencing run, yielding on the order of 10^4^–10^5^ individual sequences per sample (assuming the run is performed on an Illumina MiSeq instrument).

The level of coverage granted by a set number of reads depends on the length of the DNA sample being sequenced, the length of the NGS read used to sequence it, and whether those reads are paired-end. NGS reads are short (300 bp or less on Illumina systems), and so reads must be spread across a longer sample to sequence it in full. The expected coverage (average coverage per nucleotide) obtained for a DNA sample thus depends both on its length and the read length used for sequencing. For example, with the ~10^5^ reads returned by a commercial MiSeq multiplexed sequencing run, a 3 Mb genome could be sequenced with 150 bp paired-end reads to an expected coverage of ~10x, whereas a 20 kb plasmid could be sequenced to an expected coverage of ~1500x.

Because shorter samples can be sequenced at higher coverage for a given number of reads, it is advantageous to sequence only the region of interest of a sample. This is exemplified by amplicon sequencing, a strategy in which a researcher sequences a PCR product (an amplicon) that targets a specific region of interest in the DNA.^26^ For instance, continuing the example from above, with ~10^5^ total 150 bp paired-end reads, a 300 bp PCR product could be sequenced to an expected coverage of ~100,000x.

Many mutagenesis methods employed in protein engineering (e.g., site-saturation^27^ and tile-based mutagenesis^28^ strategies) target mutations to a specific position or region in the sequence of a protein, and thus the variants produced can be sequenced with amplicon sequencing to high coverage.^20^ Notably, however, even though increasing coverage yields more confident results, it comes with diminishing returns, and it is generally held that coverage in the tens is more than sufficient for effective reference-based identification of mutations (Supplemental Figure S1).^29^ Indeed, clinical sequencing of human genomes targets 30x coverage or greater to minimize false base calls. Given this reference, it is clear that the ~100,000x coverage that would be returned from a multiplexed sequencing run for a 300 bp amplicon is far higher than necessary for effective identification of mutations—2,000 amplicons could be sequenced in the same run and still yield clinical-grade coverage.

evSeq thus resulted from the realization that (1) at tens of dollars per sample, the cost of sending a single sample to an outsourced multiplexed NGS run is comparable to the total cost of Sanger sequencing the top variants in a round of DE, (2) amplicon sequencing can be used to identify mutations in protein variants from many protein engineering library types, and (3) enough reads are returned for a single sample in a commercial multiplexed NGS run to sequence hundreds of amplicons. Specifically, the evSeq protocol (Figure 1, *Supplemental: evSeq Library Preparation/Data Analysis Protocol*) works by focusing all reads of a single multiplexed NGS sample to specific regions on hundreds of protein variants, achieving sequencing depths of 10^1^–10^3^ at the approximate cost of existing methods for sequencing just the top variants in a round of DE (Supplemental Figure S1).

**Figure 1.**
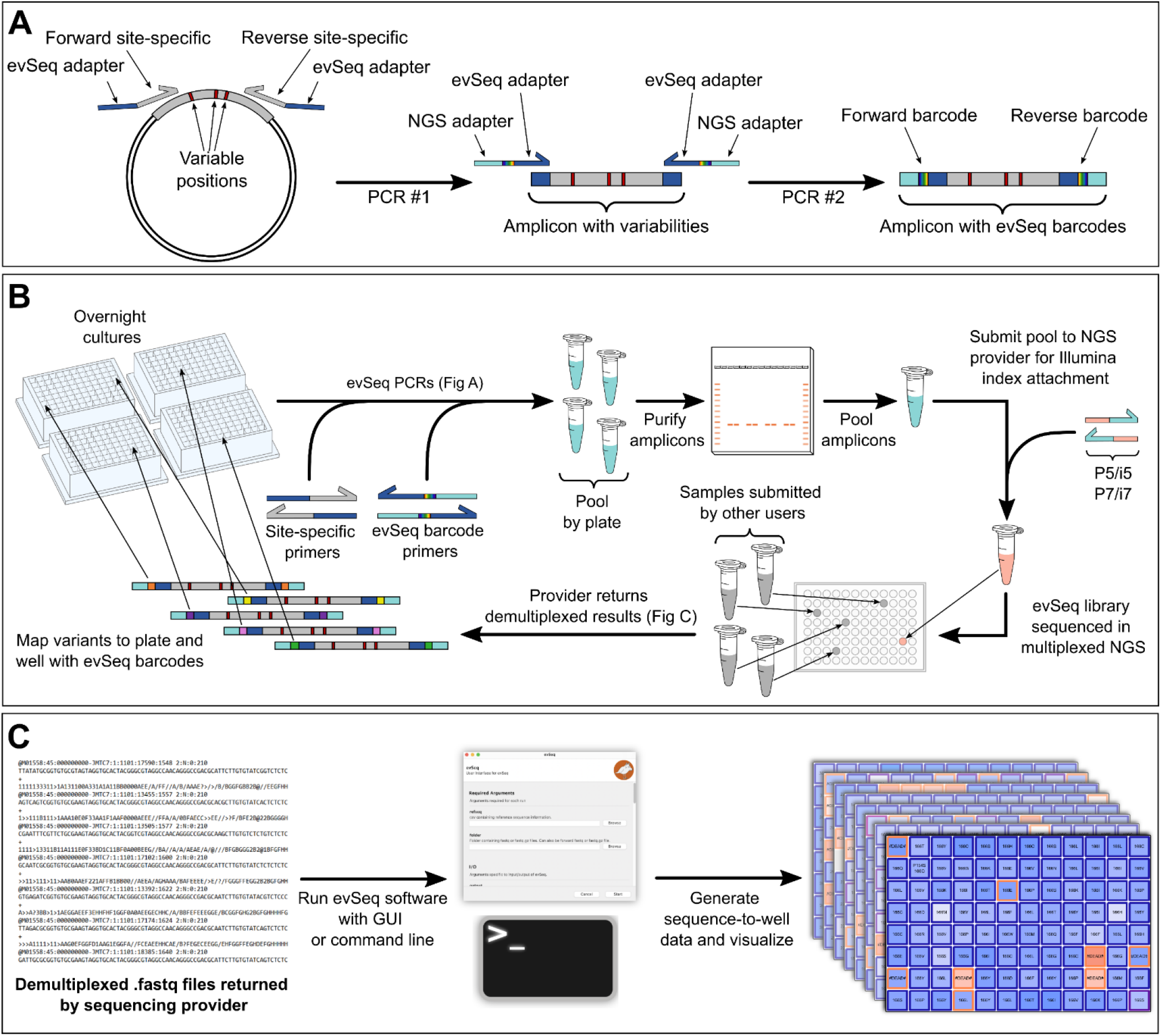
Overview of evSeq library preparation and processing. (A) In the first stage of the PCR, a region of interest is amplified with primers that include a 3’ site-specific region (gray) with 5’ adapter sequences (dark blue). The second PCR stage adds molecular barcodes (rainbow) with primers that bind to the adapter regions and add adapters for downstream NGS processing (light blue). (B) To avoid costly DNA isolation steps, evSeq uses liquid cultures of cells harboring mutated DNA (e.g., an “overnight culture” of *E. coli*) as template during the one-pot two-step barcoding PCR described in A. Each plate is pooled individually and gel purified. Purified pools are then adjusted for concentration differences and pooled together before being sent to a sequencing provider, who then appends another set of barcodes as well as sequence elements necessary for Illumina NGS sequencing. This sample is now pooled with those of other users and a multiplexed sequencing run is performed. After sequencing, the sequencing provider uses the barcodes that they attached to separate (“demultiplex”) the evSeq reads from reads of other users; the provider returns evSeq reads in .fastq files. (C) The .fastq files returned by the NGS provider are inputs to the evSeq software, which uses the evSeq forward/reverse barcode pair to map each read to a specific plate and well based on known barcode combinations. The software also processes the mapped reads (see the Supplemental and evSeq documentation for more details) to, among other things, assign variant identities to each well and return interactive HTML visualizations.

The evSeq library preparation protocol begins with PCR amplification of the region of interest in each variant (i.e., the position/region where mutations were made) and appending inline DNA barcodes to the resultant amplicons that encode their original plate-well position (Figure 1A). This is a one-pot, two-step, plate-based PCR procedure that uses two sets of primer pairs. Each primer in the first set of primers (“inner” primers) consists of a user-specified 3’ “seed” region that binds to the regions flanking the region of interest as well as a 5’ predefined universal adapter (*Supplemental: Inner Primer Design*). Each primer in the second set of primers (“outer” primers) consists of (1) a 3’ region that matches the adapter of the inner primers, (2) a central 7-nucleotide barcode where each barcode pair between forward and reverse outer primers is unique to a plate-well position, and (3) a 5’ sequence matching the Illumina Nextera transposase adapters (*Supplemental: Outer Primer Design, Supplemental: Barcode Design*, Supplemental Tables S1–S2). We designed 96 unique forward and 96 unique reverse outer primers for evSeq which, because both forward and reverse outer primers contain a barcode, can be combined to encode up to 96^2^ = 9,216 possible plate-well positions (*Supplemental: Preparation of evSeq Barcode Primer Mixes,* Supplemental Tables S3–S10. Note that we also provide a pre-filled IDT order form for the outer primers on the GitHub associated with this work—see *Supplemental: Ordering Barcode Primers from IDT* for details. While we recommend using these pre-tested barcodes, users can also design their own to, e.g., further expand the number of available combinations). Importantly, this set of outer primers can be used to sequence any target region from any gene with evSeq, and so only needs to be ordered once; only a new inner primer pair is needed for each new region of interest.

Once all barcoded amplicons have been produced, they are pooled and sent to a sequencing provider, who will then use the transposase adapters installed with the outer primers as a handle to barcode the *pool* of amplicons *once again* with a pair of sample-specific Illumina indices (Figure 1B). At this point each amplicon in the pool has one pair of sample-specific Illumina barcodes, a forward plate-well-specific inline barcode, and a reverse plate-well-specific inline barcode. This complete evSeq library is sequenced as a single sample in a multiplexed NGS run along with samples from other users (whether or not they are also evSeq samples). Post sequencing, the sequencing provider uses the sample-specific barcodes to identify those sequences belonging to the evSeq pool and returns them to the user (i.e., the provider “demultiplexes” the run, separating evSeq sequences from those of other users). The user then uses the evSeq software to analyze the returned sequences, assigning them to corresponding plate-well positions using the evSeq barcodes and identifying the mutations in the variants relative to a reference (Figure 1B, 1C).

### evSeq library preparation fits into existing protein engineering workflows and was designed to be resource efficient

A typical procedure for evaluating protein variants involves (1) arraying colonies of an organism (e.g., *Escherichia coli*) that harbor a plasmid encoding a protein variant into the wells of a (usually 96-well) microplate, (2) growing the resulting cultures to stationary phase (colloquially, an “overnight culture”), (3) using the overnight culture to inoculate a fresh culture that will be used to express the protein variants, and (4) evaluating the fitnesses of expressed protein variants. The expression stage (step 3) typically involves downtime where the experimentalist must wait until the culture reaches sufficient density before inducing protein expression and then again as expression takes place. evSeq library preparation can be performed easily in either of these time windows.

The evSeq library preparation protocol begins with the barcoding PCR described at the end of the previous section; this one-pot, two-step, plate-based PCR was designed to minimize both preparation time and laboratory resource usage (*Supplemental: evSeq Library Preparation/Data Analysis Protocol*). For instance, because evSeq relies on 96 unique forward barcodes and 96 unique reverse barcodes, a single-primer PCR would require ordering 192 new barcoding primers for each new target region evaluated in each library. In our two-primer protocol, however, the inclusion of a universal adapter on the inner primers allows the same 192 outer primers to be used regardless of target position in the variant—only two unique primers (forward and reverse inner) must be purchased for each new target region, and only if existing inner primers from previously targeted regions are not already compatible. Additionally, the evSeq PCR directly uses liquid from the overnight culture as a source of template DNA (Figure 1B, *Supplemental: evSeq Library Preparation/Data Analysis Protocol*); the template DNA is released from lysed cells during the initial heating step of the PCR, avoiding a costly and time-intensive DNA isolation/purification step and allowing researchers to use materials already prepared as part of the protein expression workflow.

The remaining steps of evSeq library preparation were, like the PCR stage, also designed to be resource and time efficient. After completion of the PCR, the resulting barcoded amplicons are pooled by plate and purified via gel extraction. Pooling prior to purification goes against standard practice for multiplexed NGS library preparation, which is to purify samples individually, quantify their DNA concentration, then combine them in equimolar quantities to ensure more equal read distribution across samples after sequencing.^30^ However, because individual plates in protein engineering libraries tend to contain variants from the same region of the same protein scaffold (e.g., as would be typical for variants from a comprehensive site-saturation library), evSeq assumes that the variation in PCR reaction yield will be minor within plates and that, as a result, the same plate can be pooled prior to quantification with only minor effects on read distribution. Using this “pooling first” strategy, only as many purifications as there are *plates* must be performed as opposed to as many as there are *variants*, thus enabling faster processing of evSeq amplicons while reducing resource usage. As will be shown in later sections, the distribution of reads returned using pooling first is perfectly acceptable for confidently identifying variant sequences.

Once all pooled plates have been purified, the concentrations of the individual purified pools are measured. The pools are then normalized by molarity and combined into a final evSeq library, which is in turn submitted as a single sample to a sequencing provider. As described in the previous section, the provider will perform a final PCR on the evSeq library to add sample-specific barcodes before sequencing it as a single sample in a multiplexed sequencing run. Outsourcing the sequencing stage has two main benefits: First, it allows evSeq to be performed by research groups with no prior sequencing experience and no direct access to sequencing equipment—groups need only be familiar with PCR, a ubiquitous technology in protein engineering laboratories. Second, to be cost effective, multiplexed sequencing should be run with tens of samples at least (Supplemental Figure S1). By outsourcing the sequencing stage, groups that do not frequently produce evSeq libraries need not wait until enough libraries have accumulated to run sequencing—a single outsourced submission, for instance, can be run along with those of other research groups with a variety of different sequencing needs.

The final stage of the evSeq workflow is data analysis using the evSeq software (github.com/fhalab/evSeq) (Figure 1C). Extensive documentation of the software and its capabilities is available as a website (fhalab.github.io/evSeq). The software was designed to be accessible to users with varying degrees of computational experience and can be run through either a graphical user interface (GUI), a command line application, or in a Python environment (e.g., a Jupyter notebook). Outputs from the software range from high-level overviews of data (e.g., an interactive “Platemap” graphic that displays sequencing coverage and identified mutations in each well of each plate; see Figure 1C for an example) to low-level details about the population of reads assigned to each well (e.g., in a well identified as polyclonal, the percentage of reads mapping to each of the identified variants). Functional data can also be easily associated with identified variants using the evSeq software outputs to produce sequence-fitness datasets, and we provide Jupyter notebooks and web pages that walk users through the process.

### evSeq facilitates library construction, validation, and sequence-fitness pairing

To highlight the utility of evSeq for engineering and biochemical experiments, we first examined how it could be used to construct high-confidence and informative sequence-fitness data. Specifically, we constructed and screened eight single site-saturation libraries of the enzyme Tm9D8*—an engineered β-subunit of tryptophan synthase from *Thermotoga maritima* (*Tm*TrpB)— for tryptophan-forming activity at 30 °C (Figure 2).^31^ In two of the screened libraries, we targeted two positions distant from the active site (A118 and S292) that have been seen to play a role in allosteric regulation of *Tm*TrpB enzymes; in the other six libraries, we targeted active-site residues known to modulate the activity of TrpB (E105, L162, I166, F184, S228, and Y301) (Figure 2A).^32–34^ As we show below, this type of sequence-fitness data can be used to assess the quality of a protein engineering library, identify improved variants during a round of directed evolution, and give insight into the significance of a given residue in catalysis.

**Figure 2.**
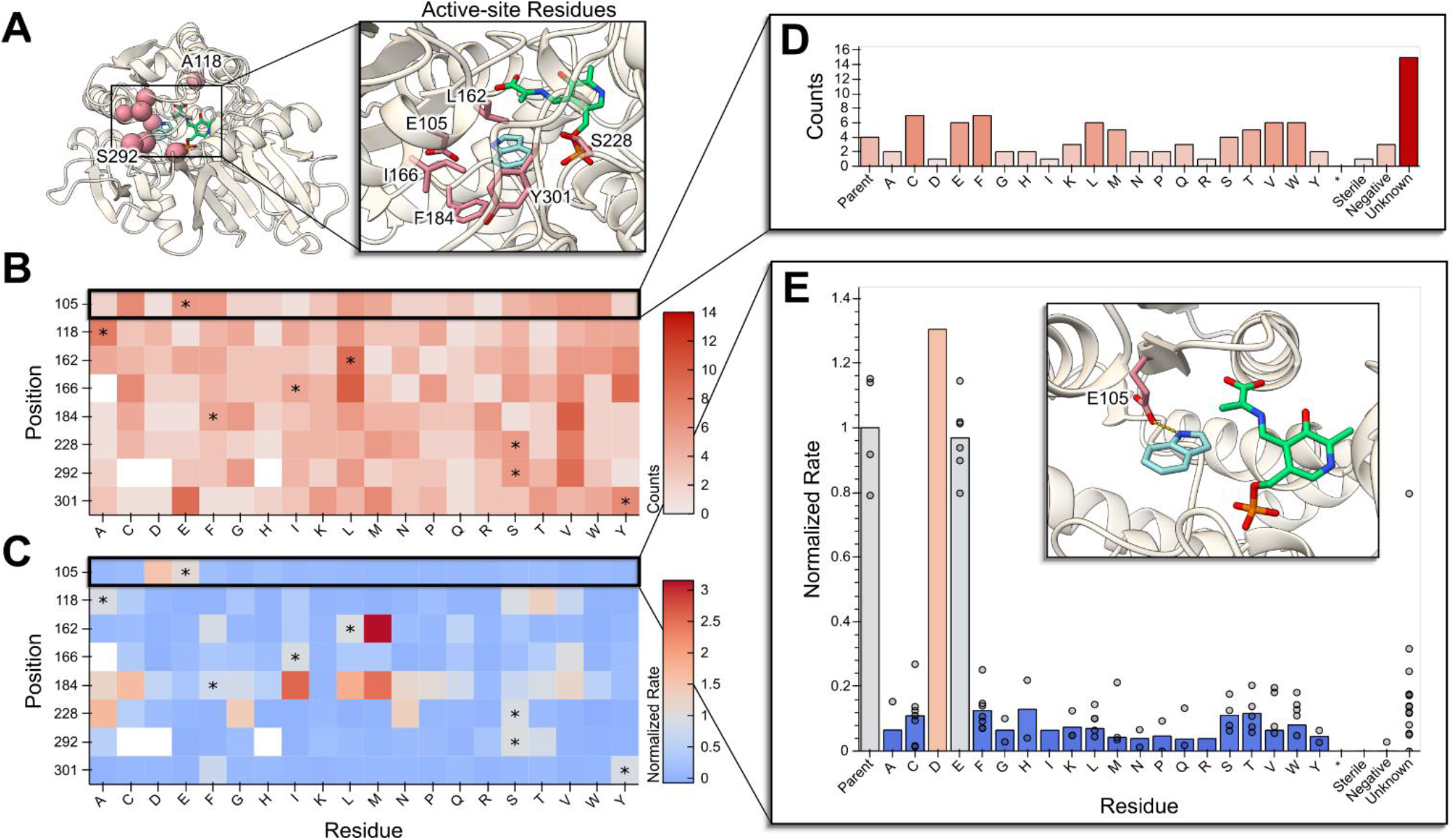
evSeq enables low-cost investigation of library quality and sequence-fitness pairing in site-saturation mutagenesis libraries. (A) Eight residues (red) known to modulate the activity of Tm9D8* were independently targeted with site-saturation mutagenesis: A118 and S292 (distal residues), E105, L162, I166, F184, S228, and Y301 (active-site residues). An active form of the pyridoxal 5’-phosphate cofactor is represented in green, and the substrate indole is shown in light blue. (B) Library quality can be investigated by plotting a heatmap of the number of times (“Counts”) each variant/mutant at each targeted position was identified from processed evSeq data. Parent amino acids are each marked with an asterisk. (C) Likewise, the effect of mutations and mutational “hotspots” can be identified by plotting a heatmap of the average activity (“Normalized Rate”) for each variant/mutation in each library when fitness data is combined with evSeq data. (D) An example plot made possible by evSeq visualization functions shows the number of times each amino acid was found in a single TrpB library (position 105), also accounting for known controls and unidentified wells. (E) Another example output of the evSeq software shows activity for a single library (position 105), showing biological replicates. The inset displays the role of the mutated residue in this library, which is to coordinate the nitrogen of the indole substrate.

Many factors can introduce bias into a site-saturation mutagenesis experiment, such as annealing bias for the native nucleotides during the PCR for library construction or contamination with the template plasmid during transformation. Without sequencing all of the variants, it is impossible to know that the library is representative of the experimental design. Since evSeq reports exactly which variants are contained in a library, researchers can leverage this to implement important quality control practices as part of the standard protein screening workflow. For instance, of all 153 possible unique variants in our eight single-site saturation libraries, we observed 149 of them (Figures 2B and C); only I166A, S292C, S292D, and S292H could not be assigned with confidence, where we define >80% abundance in a well with >10 reads as our confidence threshold. Of the variants identified, many were found in replicate (Figure 2D) due to oversampling during colony picking, which ensures that all protein variants have a chance to be found and screened (All libraries were constructed with the 22-codon trick^35^ and 88 individual colonies were screened for each library, so we expected a 98% probability of seeing all variants assuming perfect construction of libraries). Conveniently, this oversampling also allows us to evaluate the noise in our functional screen (Figure 2E) which further improves the confidence in the quality of data gathered.

Given just the fitness data gathered in this experiment, a protein engineer would identify 50 wells that are at least 1.2-fold improved over the parent enzyme Tm9D8*. However, with the sequence-fitness pairs constructed via evSeq, we know that these 50 *wells* correspond to only 16 unique *variants*. Depending on how conservative the engineer was as to what should be sequenced, this could result in anywhere from 12 (2-fold improvement) to 50 (1.2-fold improvement) wells sent off for Sanger sequencing for a total cost of $36 to $150 (using an estimate of $3 per sequence). It would cost ~$2000 to sequence all eight plates via Sanger. Using evSeq, we obtained the sequences of 625 wells of variants for only $100, corresponding to a total cost of $0.13 per non-control well, showing that evSeq is much more cost efficient for gathering sequence-fitness data from targeted mutagenesis libraries. Importantly, although the evSeq defaults currently allow only eight plates to be sequenced at once, the number of variants included in this experiment could likely have been increased as the median number of reads per well was 86 (mean: 98), which is above what is needed for reliable sequencing. Assuming that doubling the number of plates would halve the number of reads seen for each well, doubling the number of plates sequenced would cause only 14 non-control well sequences to drop below the confidence threshold.

The per-variant cost of evSeq may be reduced even further using different services and sequencing platforms. For instance, in both this section and the next, the reported number of reads and ~$100 total cost are from outsourced MiSeq runs, which returned hundreds of thousands of total reads per evSeq library. We report these numbers because outsourced multiplexed MiSeq is a standard service available to all research groups. As an alternative to outsourcing, however, our institution provides multiplexed sequencing (via the Caltech Millard and Muriel Jacobs Genetics and Genomics Laboratory) on an Illumina NextSeq platform, returning an average of ~10x more reads than the outsourced MiSeq run for a total cost of ~$10. At 10x more reads and 10x less the total cost, the per-variant evSeq cost could decrease 100-fold to <$0.01. Indeed, we were able to re-sequence the TrpB libraries at a per-variant cost of ~$0.01 with ~2.2 million total reads returned for an average of thousands of reads per variant, far higher than what is needed for reliable variant calling. It must be noted, however, that analysis of the millions of evSeq reads was no longer practical on a personal laptop, requiring a desktop workstation instead. Computational power beyond a laptop will be needed when processing more than hundreds of thousands of reads with the existing evSeq software.

Of final note, aside from providing valuable information for protein engineering experiments, evSeq can also facilitate investigation into the biochemical relevance of specific positions/mutations. Specifically, because all possible variants in a site-saturation library can be identified by evSeq, the sequence-fitness information generated can be used to explore the effects of mutations more fully than, for instance, an alanine scanning experiment.^36^ Using an example from the TrpB data gathered here, an alanine scanning experiment would tell a biochemist that the mutation to the conserved catalytic residue E105A inactivates the enzyme, with no information about the effects of other amino acid changes at this position. Using site-saturation with evSeq, we instead find that all mutations to E105 except for E105D inactivate the enzyme. The fact that glutamate and aspartate are the only amino acids containing a carboxylic acid suggests that this functional group is critical for activity (Figure 2E, with inset).

### evSeq can be used to generate data for machine learning-assisted protein engineering

We next wanted to demonstrate the utility of evSeq for advancing and applying machine learning-assisted protein engineering (MLPE). In MLPE, models are trained to learn a function that relates protein sequence to protein fitness (i.e., they learn `f(sequence) = fitness)`.^5,6,9–11^ These models are then used for rapid, low-cost *in silico* prediction of protein fitness, avoiding or greatly reducing the need for often-costly laboratory screening of variants (Figure 3).

**Figure 3.**
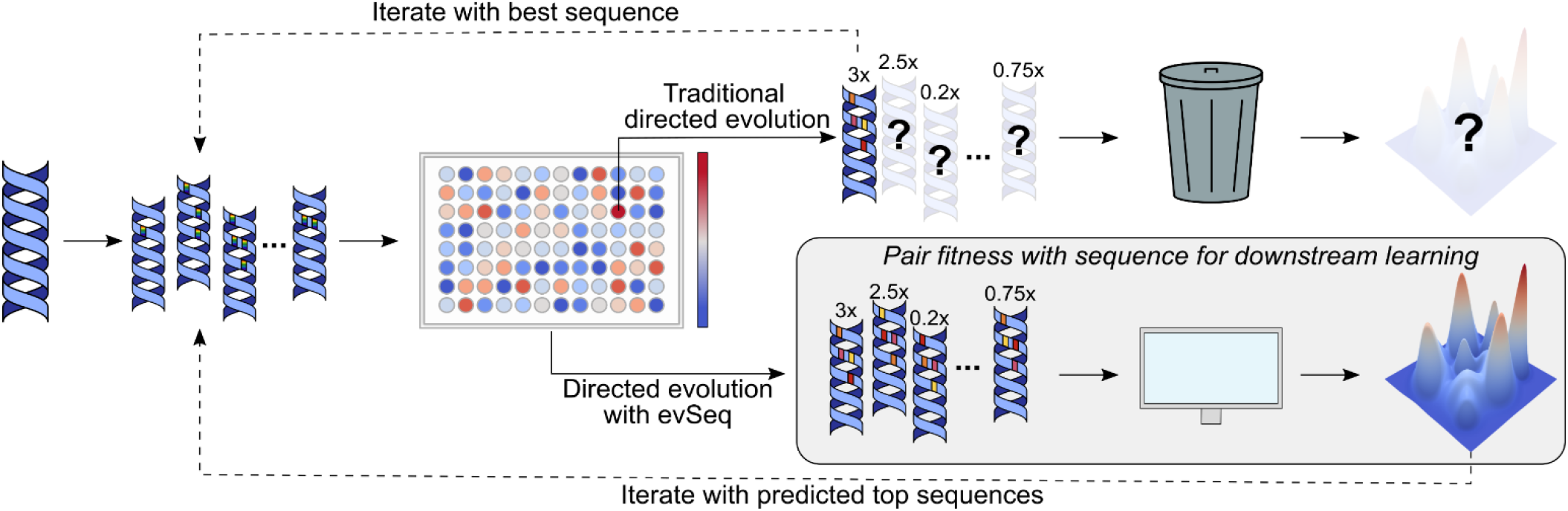
evSeq eliminates the sequencing burden of MLPE. Traditional DE only collects sequence information for top variants, essentially “throwing away” fitness data from inferior variants and learning nothing about the underlying fitness landscape. If, instead, evSeq is used to collect sequence information for all variants, MLPE methods, which require sequence-fitness pairs for supervised model training, can be implemented. Sampling from a fitness landscape, an ML model can be trained to predict the fitnesses of missing sequences and reconstruct the missing regions of this landscape.

Sequence-fitness data is critical for effective MLPE. Indeed, even though strategies exist that *can* predict protein fitness from sequence alone (e.g., those that use evolutionary data to predict protein fitness), their effectiveness is improved with the inclusion of sequence-fitness information.^7,14,15,37^ As a result, the most effective MLPE workflows require that both sequence *and* fitness data be collected, unlike a DE workflow, which requires only fitness data.

The need to collect sequence data in addition to fitness data is an often-overlooked additional cost of MLPE strategies compared to standard DE. For instance, we recently developed an ML strategy known as machine learning-assisted directed evolution (MLDE) for efficient navigation of epistatic combinatorial protein variant libraries.^38,39^ Previously, we used MLDE to evolve *Rhodothermus marinus* nitric oxide dioxygenase (*Rma*NOD) for greater enantioselectivity in a carbon–silicon bond-forming reaction.^39^ Over the course of the engineering campaign, we collected six 96-well plates of sequence-fitness data for training ML models. In total, sequencing the variants in these plates by Sanger sequencing cost ~$1700. High additional sequencing costs like these can make MLPE methods far less attractive, even if they are more effective than traditional DE at finding high-fitness protein variants.^38^ However, given that evSeq enables sequencing all variants for a cost similar to standard DE methods, it enables use of MLPE without added cost. In essence, evSeq eliminates the sequencing burden of MLPE.

To demonstrate the application of evSeq to MLPE, we used it to sequence five plates of *Rma*NOD variants from a four-site combinatorial library. Coupled with fitness data, the sequences resulting from this run could be used to drive a round of MLDE. Notably, sequencing these plates by Sanger sequencing would have cost ~$1400; in contrast, sequencing by evSeq using an outsourced multiplexed MiSeq run cost ~$100 for a per-variant cost of ~$0.21. The median read depth per variant in this run was 463 (mean: 506), much higher than is required for accurate sequencing, and so more plates—from either the same or a different library—could have reasonably been added to this evSeq run to decrease the per-variant sequencing cost even further (Figure 3B). Of course, as discussed in the previous section, in-house sequencing could have cut sequencing costs an additional ten-fold.

The cost of sequencing is most notably a barrier for MLPE strategies that focus on developing models for a single protein with a well-defined fitness (e.g., MLDE); however, the applicability of evSeq to MLPE is not limited solely to cost-reduction. For instance, ML strategies have been developed that, rather than focusing on a specific protein, train models on sequence-fitness information across multiple different protein scaffolds.^16,40^ The goal is for these models to learn global determinants of protein fitness, then to use the models as general-purpose protein fitness predictors. By enabling the collection of sequence-fitness pairs across a wider array of proteins and fitness definitions, evSeq opens these approaches to new and more diverse data sources. Generally speaking, the more sequence-fitness data available to train and benchmark these strategies, the better we expect them to perform and the more rapidly we expect improvements to be developed.^16^ It is no coincidence that large leaps forward in other ML disciplines have followed increased availability of large, diverse datasets, with the rapid advance in computer vision sparked by ImageNet being perhaps the most prominent example.^41^ Widespread adoption of evSeq—and commitment to depositing sequence-fitness data in resources such as ProtaBank—would provide such a dataset for protein engineering. This dataset would span the range of all engineered proteins and all target fitnesses, capture examples of sequences with both higher and lower/zero fitness relative to a parent (the latter of which is effectively never recorded with current DE sequencing practices), and overall enable rapid advancement in MLPE.^8^

### evSeq detects all variability in the sequenced amplicons

Although we focused here on demonstrating applications involving targeted mutagenesis strategies, evSeq is also applicable to other mutagenesis methods, as the associated software can identify both user-specified and unspecified positions of variability (Figure 4A). This feature not only informs the user of potential unexpected mutations in the sequenced amplicon (Supplemental Table S11), but also allows it to work effectively with tile-based mutagenesis strategies and other semi-targeted mutagenesis strategies (e.g., error-prone PCR of specific regions or small genes). All that is required is that the amplicon length and read length be able to capture the full region containing mutations.

**Figure 4.**
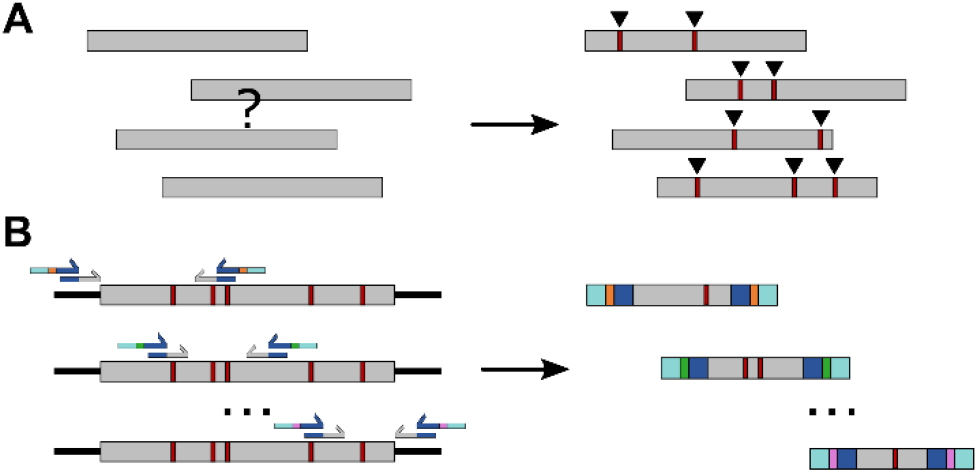
evSeq detects variability and can be expanded for random mutagenesis. (A) evSeq does not require that the user specify which position in the amplicon was targeted. Instead, the software can identify variable regions by comparing to a reference (B) evSeq can be used to sequence entire genes by designing a set of inner primer pairs which together capture the entire gene. Different evSeq barcodes can then be used for each region, and the user can reconstruct the entire sequence.

It should be noted that evSeq will not detect off-target mutations outside of the constructed amplicon as these regions are not sequenced, meaning that it is unable to identify other mutations in a larger DNA element that may be contributing to activity. Due to this fact, for exceedingly unexpected mutational effects that are not seen in replicate, we suggest sequencing the rest of the DNA element to confirm the presence or absence of any off-target mutations. However, this limitation is mitigated by the fact that off-target mutations are rare and, importantly, evSeq is agnostic to read length and will work with any length of paired-end sequencing. While the current software version is not yet suited for other, long-read sequencing technologies (e.g., PacBio or Oxford Nanopore), future versions could be updated and validated with these data formats and make full gene-length evSeq experiments more straightforward and cost effective. Given this, evSeq is currently best suited and most cost effective when all expected mutations exist in the sequenced amplicon, though sequencing of multiple overlapping amplicons can readily allow evSeq to be expanded to sequence entire genes of variants arrayed in microplates (Figure 4B).

## CONCLUSION

Hundreds to thousands of protein variants (or more) are constructed and their fitnesses evaluated over the course of a standard protein engineering campaign. Without sequencing, these fitnesses are next to useless—the time, effort, and resources expended to produce them are largely wasted. As a strategy that is comparable in cost to existing protocols, accessible to non-computational scientists, and easy to implement with existing technology, evSeq rescues these fitness data by making the collection of sequence data for every variant a practical and highly useful step of the protein engineering pipeline. Given the number of research groups working on DE and other protein engineering projects, widespread adoption of this technology would lead to an explosion in the availability of sequence-fitness information. By sequencing every variant, no laboratory screening effort is wasted, and we open the door to advances in both our biochemical understanding of proteins and our ability to engineer them with data-driven methods.

## AUTHOR INFORMATION

### Corresponding Author

Frances H. Arnold – Division of Biology and Biological Engineering, Division of Chemistry and Chemical Engineering, California Institute of Technology, MC 210-41, 1200 E. California Boulevard, Pasadena, CA 91125, USA;

### Authors

Bruce J. Wittmann – Division of Biology and Biological Engineering, California Institute of Technology, MC 210-41, 1200 E. California Boulevard, Pasadena, CA 91125, USA;

Kadina E. Johnston – Division of Biology and Biological Engineering, California Institute of Technology, MC 210-41, 1200 E. California Boulevard, Pasadena, CA 91125, USA;

Patrick J. Almhjell – Division of Chemistry and Chemical Engineering, California Institute of Technology, MC 210-41, 1200 E. C lifornia Boulevard, Pasadena, CA 91125, USA;

### Author Contributions

Author contributions are provided using the CRediT taxonomy: **BJW:** Conceptualization, methodology, software, validation, investigation, writing – original draft, writing – review and editing, funding acquisition. **KEJ:** Methodology, data-collection, software, investigation, writing – original draft, writing – review and editing, visualization, funding acquisition. **PJA:** Methodology, data-collection, software, validation, investigation, writing – original draft, writing – review and editing, visualization. **FHA:** Resources, writing – original draft, writing – review and editing, funding acquisition.

## ACKNOWLEDGMENTS

The authors thank Shan Li, Adrienne Rollie, and Eric Brustad at Illumina, Inc: Shan Li and Adrienne Rollie for helping us troubleshoot the evSeq method and Eric Brustad for critical reading of the manuscript. We also thank fellow Arnold laboratory members Nathaniel Goldberg and Nicholas Porter for implementing evSeq (which pointed us to necessary improvements in the method), Anders Knight for suggesting and prototyping evSeq software features, and Sabine Brinkmann-Chen for critical reading of the manuscript.

## ABBREVIATIONS

evSeq: every variant sequencing
DE: directed evolution
ML: machine learning
DMS: deep mutational scanning
NGS: next-generation sequencing
GUI: graphic user interface
*Tm*TrpB: *Thermotoga maritima* β-subunit of tryptophan synthase
MLPE: machine learning-assisted protein engineering
MLDE: machine learning-assisted directed evolution
*Rma*NOD: *Rhodothermus marinus* nitric oxide dioxygenase

## REFERENCES

(1) BCC Research Staff: Global Markets for Enzymes in Industrial Applications. BCC Research LLC; 2018

(2) Podgornaia, A. I.; Laub, M. T. Pervasive Degeneracy and Epistasis in a Protein-Protein Interface. Science. 2015, 347 (6222), 673–677.

(3) Wu, N. C.; Dai, L.; Olson, C. A.; Lloyd-Smith, J. O.; Sun, R. Adaptation in Protein Fitness Landscapes Is Facilitated by Indirect Paths. Elife 2016, 5, e16965.

(4) Faure, A. J.; Domingo, J.; Schmiedel, J. M.; Hidalgo-Carcedo, C.; Diss, G.; Lehner, B. Global Mapping of the Energetic and Allosteric Landscapes of Protein Binding Domains. bioRxiv 2021, 1–45.

(5) Yang, K. K.; Wu, Z.; Arnold, F. H. Machine-Learning-Guided Directed Evolution for Protein Engineering. Nat. Methods 2019, 16 (8), 687–694.

(6) Li, G.; Dong, Y.; Reetz, M. T. Can Machine Learning Revolutionize Directed Evolution of Selective Enzymes? Adv. Synth. Catal. 2019, 361 (11), 2377–2386.

(7) Hopf, T. A.; Ingraham, J. B.; Poelwijk, F. J.; Schärfe, C. P. I.; Springer, M.; Sander, C.; Marks, D. S. Mutation Effects Predicted from Sequence Co-Variation. Nat. Biotechnol. 2017, 35 (2), 128–135.

(8) Wang, C. Y.; Chang, P. M.; Ary, M. L.; Allen, B. D.; Chica, R. A.; Mayo, S. L.; Olafson, B. D. ProtaBank: A Repository for Protein Design and Engineering Data. Protein Sci. 2018, 27 (6), 1113–1124.

(9) Mazurenko, S.; Prokop, Z.; Damborsky, J. Machine Learning in Enzyme Engineering. ACS Catal. 2020, 10 (2), 1210–1223.

(10) Siedhoff, N. E.; Schwaneberg, U.; Davari, M. D. Machine Learning-Assisted Enzyme Engineering. In Methods in Enzymology; Elsevier Inc., 2020; Vol. 643, pp 281–315.

(11) Wittmann, B. J.; Johnston, K. E.; Wu, Z.; Arnold, F. H. Advances in Machine Learning for Directed Evolution. Curr. Opin. Struct. Biol. 2021, 69, 11–18.

(12) Fowler, D. M.; Fields, S. Deep Mutational Scanning: A New Style of Protein Science. Nat. Methods 2014, 11 (8), 801–807.

(13) Livesey, B. J.; Marsh, J. A. Using Deep Mutational Scanning to Benchmark Variant Effect Predictors and Identify Disease Mutations. Mol. Syst. Biol. 2020, 16 (7), e9380.

(14) Riesselman, A. J.; Ingraham, J. B.; Marks, D. S. Deep Generative Models of Genetic Variation Capture the Effects of Mutations. Nat. Methods 2018, 15 (10), 816–822.

(15) Meier, J.; Rao, R.; Verkuil, R.; Liu, J.; Sercu, T.; Rives, A. Language Models Enable Zero-Shot Prediction of the Effects of Mutations on Protein Function. bioRxiv 2021, 1–28.

(16) Gray, V. E.; Hause, R. J.; Luebeck, J.; Shendure, J.; Fowler, D. M. Quantitative Missense Variant Effect Prediction Using Large-Scale Mutagenesis Data. Cell Syst. 2018, 6 (1), 116–124.

(17) Sanger, F.; Coulson, A. R. A Rapid Method for Determining Sequences in DNA by Primed Synthesis with DNA Polymerase. J. Mol. Bid 1975, 94 (3), 441–448.

(18) Metzker, M. L. Sequencing Technologies — the next Generation. Nat. Rev. Genet. 2010, 11 (1), 31–46.

(19) Smith, A. M.; Heisler, L. E.; St.Onge, R. P.; Farias-Hesson, E.; Wallace, I. M.; Bodeau, J.; Harris, A. N.; Perry, K. M.; Giaever, G.; Pourmand, N.; et al. Highly-Multiplexed Barcode Sequencing: An Efficient Method for Parallel Analysis of Pooled Samples. Nucleic Acids Res. 2010, 38 (13), e142.

(20) Appel, M. J.; Longwell, S. A.; Morri, M.; Neff, N.; Herschlag, D.; Fordyce, P. M. UPIC–M: Efficient and Scalable Preparation of Clonal Single Mutant Libraries for High-Throughput Protein Biochemistry. bioRxiv 2021, 1–18.

(21) Srivathsan, A.; Lee, L.; Katoh, K.; Hartop, E.; Kutty, S. N.; Wong, J.; Yeo, D.; Meier, R. MinION Barcodes: Biodiversity Discovery and Identification by Everyone, for Everyone. bioRxiv 2021, 1–52.

(22) Glenn, T. C.; Nilsen, R. A.; Kieran, T. J.; Sanders, J. G.; Bayona-Vásquez, N. J.; Finger, J. W.; Pierson, T. W.; Bentley, K. E.; Hoffberg, S. L.; Louha, S.; et al. Adapterama I: Universal Stubs and Primers for 384 Unique Dual-Indexed or 147,456 Combinatorially-Indexed Illumina Libraries (ITru & INext). PeerJ 2019, 7, e7755.

(23) Wierbowski, S. D.; Vo, T. V; Falter-Braun, P.; Jobe, T. O.; Kruse, L. H.; Wei, X.; Liang, J.; Meyer, M. J.; Akturk, N.; Rivera-Erick, C. A.; et al. A Massively Parallel Barcoded Sequencing Pipeline Enables Generation of the First ORFeome and Interactome Map for Rice. Proc. Natl. Acad. Sci. 2020, 117 (21), 11836–11842.

(24) Chubiz, L. M.; Lee, M.-C.; Delaney, N. F.; Marx, C. J. FREQ-Seq: A Rapid, Cost-Effective, Sequencing-Based Method to Determine Allele Frequencies Directly from Mixed Populations. PLoS One 2012, 7 (10), e47959.

(25) Weile, J.; Sun, S.; Cote, A. G.; Knapp, J.; Verby, M.; Mellor, J. C.; Wu, Y.; Pons, C.; Wong, C.; van Lieshout, N.; et al. A Framework for Exhaustively Mapping Functional Missense Variants. Mol. Syst. Biol. 2017, 13 (12), 957.

(26) Wen, C.; Wu, L.; Qin, Y.; Van Nostrand, J. D.; Ning, D.; Sun, B.; Xue, K.; Liu, F.; Deng, Y.; Liang, Y.; et al. Evaluation of the Reproducibility of Amplicon Sequencing with Illumina MiSeq Platform. PLoS One 2017, 12 (4), e0176716.

(27) Siloto, R. M. P.; Weselake, R. J. Site Saturation Mutagenesis: Methods and Applications in Protein Engineering. Biocatal. Agric. Biotechnol. 2012, 1 (3), 181–189.

(28) Melnikov, A.; Rogov, P.; Wang, L.; Gnirke, A.; Mikkelsen, T. S. Comprehensive Mutational Scanning of a Kinase in Vivo Reveals Substrate-Dependent Fitness Landscapes. Nucleic Acids Res. 2014, 42 (14), e112.

(29) Sims, D.; Sudbery, I.; Ilott, N. E.; Heger, A.; Ponting, C. P. Sequencing Depth and Coverage: Key Considerations in Genomic Analyses. Nat. Rev. Genet. 2014, 15 (2), 121–132.

(30) Illumina. Nextera XT DNA Library Prep Reference Guide. 2019.

(31) Boville, C. E.; Romney, D. K.; Almhjell, P. J.; Sieben, M.; Arnold, F. H. Improved Synthesis of 4-Cyanotryptophan and Other Tryptophan Analogues in Aqueous Solvent Using Variants of TrpB from Thermotoga Maritima. J. Org. Chem. 2018, 83 (14), 7447–7452.

(32) Rix, G.; Watkins-Dulaney, E. J.; Almhjell, P. J.; Boville, C. E.; Arnold, F. H.; Liu, C. C. Scalable Continuous Evolution for the Generation of Diverse Enzyme Variants Encompassing Promiscuous Activities. Nat. Commun. 2020, 11 (1), 5644.

(33) Romney, D. K.; Murciano-Calles, J.; Wehrmüller, J. E.; Arnold, F. H. Unlocking Reactivity of TrpB: A General Biocatalytic Platform for Synthesis of Tryptophan Analogues. J. Am. Chem. Soc. 2017, 139 (31), 10769–10776.

(34) Buller, A. R.; Brinkmann-Chen, S.; Romney, D. K.; Herger, M.; Murciano-Calles, J.; Arnold, F. H. Directed Evolution of the Tryptophan Synthase β-Subunit for Stand-Alone Function Recapitulates Allosteric Activation. Proc. Natl. Acad. Sci. 2015, 112 (47), 14599–14604.

(35) Kille, S.; Acevedo-Rocha, C. G.; Parra, L. P.; Zhang, Z.-G.; Opperman, D. J.; Reetz, M. T.; Acevedo, J. P. Reducing Codon Redundancy and Screening Effort of Combinatorial Protein Libraries Created by Saturation Mutagenesis. ACS Synth. Biol. 2013, 2 (2), 83–92.

(36) Morrison, K. L.; Weiss, G. A. Combinatorial Alanine-Scanning. Curr. Opin. Chem. Biol. 2001, 5 (3), 302–307.

(37) Hsu, C.; Nisonoff, H.; Fannjiang, C.; Listgarten, J. Combining Evolutionary and Assay-Labelled Data for Protein Fitness Prediction. bioRxiv 2021.

(38) Wittmann, B. J.; Yue, Y.; Arnold, F. H. Informed Training Set Design Enables Efficient Machine Learning-Assisted Directed Protein Evolution. Cell Syst. 2021, 12 (11), 1026–1045.e7.

(39) Wu, Z.; Kan, S. B. J.; Lewis, R. D.; Wittmann, B. J.; Arnold, F. H. Machine Learning-Assisted Directed Protein Evolution with Combinatorial Libraries. Proc. Natl. Acad. Sci. 2019, 116 (18), 8852–8858.

(40) Alieva, A.; Aceves, A.; Song, J.; Mayo, S.; Yue, Y.; Chen, Y. Learning to Make Decisions via Submodular Regularization. ICLR 2021.

(41) Russakovsky, O.; Deng, J.; Su, H.; Krause, J.; Satheesh, S.; Ma, S.; Huang, Z.; Karpathy, A.; Khosla, A.; Bernstein, M.; et al. ImageNet Large Scale Visual Recognition Challenge. Int. J. Comput. Vis. 2015, 115 (3), 211–252.

