## Supplemental Material for "evSeq: Cost-Effective Amplicon Sequencing of Every Variant in a Protein Library"

### Data Availability

All raw and processed data generated by this study can be found at CaltechData (DOI: 10.22002/D1.2140). The software version used to analyze all data in this study is tagged as v1.0.0 on the associated GitHub repository.

### Oligo Design

#### Inner Primer Design

The inner primers of evSeq are specific to the region of interest. Each region of interest is captured by both a forward and reverse primer. These primers have the below general layout:

F: 5′ – CACCCAAGACCACTCTCCGG**XXXXXXX**… – 3′

R: 5′ – CGGTGTGCGAAGTAGGTGC**XXXXXXXX**… – 3′

The 5' region is a universal adapter to which outer primers bind (see *Preparation of evSeq Barcode Primer Mixes*, below) while the 3' region (denoted by "**X**" in the primers above) is specific to the region of interest. Note that the length of the variable 3' region will vary depending on the target gene (this is indicated by the ellipses at the end of the poly-**X** region). Note that there is no need for the two primers in the pair to be equal length—we show them as such to highlight the fact that the forward universal adapter is one base longer than the reverse universal adapter. Detailed instructions for effective primer construction are provided on the evSeq wiki (https://fhalab.github.io/evSeq/1-lib_prep.html#inner-primer-design).

#### Outer Primer Design

The barcode (outer) primers used in evSeq all follow the below layout:

F: 5' – TCGTCGGCAGCGTCAGATGTGTATAAGAGACAG**XXXXXXX**CACCCAAGACCACTCTCCGG – 3'

R: 5' – GTCTCGTGGGCTCGGAGATGTGTATAAGAGACAG**XXXXXXX**CGGTGTGCGAAGTAGGTGC – 3'

Each of these primers consists of (1) a 5' sequence matching the Illumina Nextera transposase adapters, (2) a central unique 7-nucleotide barcode (Table S1), and (3) a 3' universal seed that matches the 5' adapter of the inner primers (see *Inner Primer Design*, above). Note that only Illumina indices compatible with the Nextera transposase adapters can be used with the provided outer primer designs; other indexing systems would require different adapters. The full set of outer primers used in this study can be found in Table S2; they can be ordered from IDT by following the instructions provided in *Ordering Barcode Primers from IDT*, below.

#### Barcode Design

evSeq uses 192 unique 7-nucleotide barcodes (Table S1). The barcodes were designed to satisfy the below criteria:

1. All barcodes must have GC-content of 40–60%.
2. All barcodes must be at least 3 substitutions apart. This is to prevent misassignment of reads due to sequencing errors of the barcodes.
3. No barcode can have 3 of the same bases in a row. This is to reduce sequencing errors.
4. No barcode can be a sub-sequence of the Nextera transposase adapters or their reverse complements (see below). This is to avoid interference with downstream Illumina chemistry.
5. No barcode can be a sub-sequence of the Illumina p5 and p7 flow cell-binding sequences or their reverse complements (see below for sequences). Again, this is to avoid interference with downstream Illumina chemistry.

The Nextera transposase adapter sequences are below:

5’ – TCGTCGGCAGCGTCAGATGTGTATAAGAGACAG – 3'

5’ – GTCTCGTGGGCTCGGAGATGTGTATAAGAGACAG – 3'

The p5 and p7 flow cell-binding sequences are below:

p5: 5’ – AATGATACGGCGACCACCGAGATCTACAC – 3'

p7: 5’ – CAAGCAGAAGACGGCATACGAGAT – 3’

### Supplemental Protocols

#### Ordering Barcode Primers from IDT

We provide a pre-filled IDT order form for all evSeq primers on the evSeq GitHub repository (https://github.com/fhalab/evSeq/tree/master/lib_prep_tools/IdtOrderForm.xlsx). This order form can be used to order evSeq primers in the 96-well plate layout needed to prepare the evSeq barcode primer mixes (see *Preparation of evSeq Barcode Primer Mixes*, below). To order evSeq primers:

1. Navigate to the IDT DNA oligo ordering page: https://www.idtdna.com/pages/products/custom-dna-rna/dna-oligos/custom-dna-oligos.
2. Under “Ordering”, select “Plates”.
3. From the “Single-stranded DNA” table, select the amount (in nanomoles) of oligo you wish to order (denoted in the “Product” column) by clicking “Order” under the “96 Well” column. For the work described in this paper, 25 nmole oligos were ordered.
4. On the next page, click “UPLOAD PLATE(S)”. Using the pop-up that results, upload the “IdtOrderForm.xls” provided on the evSeq GitHub repository. The pop-up should recognize two plates—one called “FBC” and the other called “RBC”—each consisting of 96 wells. Click “ADD PLATES” followed by “CLOSE THIS WINDOW” to close the window.
5. For the “FBC” plate, click “Plate Specifications”. Confirm that the below specifications are set as follows:
    a. Purification: Standard Desalting
    b. Plate Type: Deep Well
    c. **Ship Option: Wet**
    d. Buffer: IDTE 8.0 pH
    e. Normalization Type: Full Yield
    f. **Concentration: 100 μM**

    Note that the bolded specifications are different from default. **While not strictly required, it is strongly recommended that primers be ordered *wet* at 100 μM; reconstituting plates of dry primers to 100 μM can be very time-consuming without robotic support.**
6. Once specifications are correctly set for the “FBC” plate, click “APPLY SETTINGS TO ALL PLATES” at the bottom of the specifications pop-up, followed by “YES” on the window that follows. Quickly check to make sure that the same settings as recommended in step 5 were applied to “RBC” by clicking on the “RBC” “Plate Specifications” option.
7. Add the primers to your order by clicking “ADD TO ORDER”, then follow standard IDT procedures for purchasing.

#### Preparation of evSeq Barcode Primer Mixes

There are 96 unique forward and 96 unique reverse outer primers (Table S2), corresponding to 96 unique forward and 96 unique reverse barcodes (Table S1). The forward and reverse outer primers were ordered following the procedure given above in *Ordering Barcode Primers from IDT*.

Each well sequenced in evSeq is encoded by a different combination of forward and reverse barcode. Different primers from the forward and reverse outer primer plates can be mixed together to associate a barcode combination with a specific well in a specific plate. Because the same outer primers can be used regardless of inner primer, it is convenient to keep plates of barcode combinations on hand. Plates of outer primer combinations (hereafter also referred to as “barcode plates”) can be stored for long periods of time.

Throughout this work, we used the same 8 barcode plates (consisting of 768 different combinations of forward and reverse outer primers) to encode plate and well locations. Barcode plates are named DI01–DI08, where “DI” stands for “dual-indexed”. The exact barcode combinations used by evSeq are given in Table S3 – Table S10; these combinations can also be found in the “index_map.csv” file on the evSeq GitHub: (https://github.com/fhalab/evSeq/tree/master/evSeq/util/index_map.csv). By default, the evSeq software assumes the barcode plates used for library preparation are laid out in the order given in the “index_map.csv” file. To build the barcode plates depicted in Table S3 – Table S10, we followed the below procedure:

1. 10-fold dilutions of each of the forward and reverse outer primer plates ordered from IDT were prepared by adding 10 µL of each primer stock to 90 µL ddH2O, keeping the well layout constant. Dilutions were performed in fully-skirted PCR plates (Bio-Rad HSP9601). The plates from IDT had a starting concentration of 100 µM, so the final concentration of these two diluted plates was 10 µM.
2. To 8 fully-skirted PCR plates, 80 µL ddH2O was added, followed by 10 µL diluted (10 µM) forward barcode plate. The well layout was kept constant for the forward barcode primers.
3. To the each of the 8 plates, 10 µL of diluted (10 µM) reverse barcode plate was added, shifting the well layout down by 1 row per plate. For instance, row A of the reverse plate went into row A of the first barcode plate, row B of the second barcode plate, row C of the third barcode plate, and so on; row H of the reverse plate went into row H of the first barcode plate, row A of the second barcode plate, row B of the third barcode plate, and so on.
4. When not in use, the 10-fold dilutions prepared in step 1 were stored at –20 °C, while the barcode plates (each well of which had a combination of a specific forward and reverse primer at a final concentration of 1 µM) were stored at 4 °C. Both the 10 µM stock plates and 1 µM barcode plates can be stored for long periods of time—we have noticed no drop in effectiveness even after years of storage.

#### evSeq Library Preparation/Data Analysis Protocol

The evSeq library preparation protocol was designed to be as cost-effective as possible. The quantities used in the below protocol were chosen to fit within the constraints of the resources available to our research group (these are the quantities used for all evSeq experiments performed in this paper). However, with automation support (e.g., liquid handling robots) and higher-capacity molecular biology equipment, the entire protocol could be scaled down to lower quantities, further improving cost-effectiveness.

The list of steps below can be followed to prepare an evSeq library for sequencing using the outer primers described in *Preparation of evSeq Barcode Primer Mixes*, above. Note that when first using a new set of inner primers, it is recommended to complete the below protocol for a few wells as a test before deploying them for plate-scale reactions.

The library preparation protocol can be completed with the below steps. Note that provided part numbers are for the materials/reagents we used while developing this protocol—the same components from other providers will almost certainly work as well. This protocol is also provided on the evSeq wiki (https://fhalab.github.io/evSeq/1-lib_prep.html#pcr-protocol).

1. Prepare a PCR master mix for the number of wells to be sequenced according to the below table. Note that we provide an excel calculator on the evSeq GitHub repository for easy calculation of

master mix volumes based on the number of plates to be sequenced (https://github.com/fhalab/evSeq/tree/master/lib_prep_tools/MastermixCalculator.xlsx).

| Component | Amount per 10 µL rxn (µL) |
|---|---|
| Thermopol Buffer (NEB B9004S) | 1.00 |
| 10 mM dNTPs (NEB N0447) | 0.20 |
| Taq Polymerase (NEB M0267) | 0.05 |
| $ddH_2O$ | 5.33 |
| Mol-Bio Grade DMSO (MP 194819) | 0.40 |
| Inner Primer Mix (10 µM) | 0.02 |

a. Note that the above table assumes that each evSeq PCR reaction will be 10 µL—if scaling down, adjust volumes accordingly.
b. Note that the above table also assumes the same set of inner primers is used to prepare all plates. If this is not the case, a separate master mix will need to be prepared for each set of inner primers.
c. The Inner Primer Mix (10 µM) is a combination of forward and reverse inner primers at a final concentration of 10 µM each in diH2O (this can be prepared, e.g., by adding 10 µL of 100 µM forward inner primer and 10 µL of 100 µM reverse inner primer to 80 µL diH2O).

2. Add 7 µL of master mix to each well of as many half-skirted PCR plates (USA Scientific 1402-9700) as will be sequenced. These are referred to as “PCR plates” in the remainder of this protocol.
3. Stamp 1 µL of overnight culture from each plate to be sequenced into the PCR plates.
    a. “Stamp” means “apply to all wells, keeping the plate layout consistent”. For example, 1 µL of culture from library 01 F02 is moved to PCR plate 01 F02, 1 µL of culture from library 02 C07 is moved to PCR plate 02 C07, etc.
    b. Note that both fresh culture and previously frozen culture (thawed before use as template) will work here. No modifications need to be made to the protocol.
4. Complete stage 1 PCR using the below thermal cycler conditions. This PCR amplifies the fragment of interest from the template DNA contained in the cell culture.

| Step | Temperature (°C) | Time |
|---|---|---|
| 1 | 95 | 5 min |
| 2 | 95 | 20 s |
| 3 | TD 63-> 54 | 20 s |
| 4 | 68 | 30 s |
| 5 | Return to 2, 9 x | |
| 6 | 4 | Hold |

    a. "TD” above stands for “touchdown”. A touchdown step decrements the temperature by 1 °C each cycle. The touchdown in the above PCR starts at 63 °C and drops to 54 °C by the end.
    b. Note that the extension step (step 4) is long enough to amplify a 500 bp fragment. Longer fragments will need a longer extension time. Note, however, that you may see reduced sequencing efficiency with fragments that are too large.
    c. While developing this protocol, we used the below thermal cycler models:
        i. Eppendorf Mastercycler ep Gradient S Thermal Cycler, Model 5345 with 96-well universal block
        ii. Eppendorf Mastercycler pro S vapo.protect
        iii. Eppendorf Mastercycler X50s 96-well silver block thermal cycler
5. Once PCR has completed, stamp 2 µL of 1 µM barcode primer mix from the barcode plates into the PCR plates (see *Preparation of evSeq Barcode Primer Mixes*, above, for details on preparation of barcode plates). **Record which barcode plate was stamped into which PCR plate.**
6. Perform the second step PCR using the below conditions:

| Step | Temperature (°C) | Time |
|---|---|---|
| 1 | 95 | 20s |
| 2 | 68 | 50 s |
| 3 | Return to 1, 24 x | |
| 4 | 68 | 5 min |
| 5 | 4 | Hold |

   a. Again, longer fragments may need a longer extension time.

7. While the second PCR runs, prepare a 2% agarose gel with SYBR gold added (Thermo Fisher Scientific, S11494).
8. Once the second PCR has completed, add 5 µL of each reaction to 1 µL 100 mM EDTA to quench the reactions. For each plate, pool the quenched reactions. Pooling will leave you with as many tubes as you have plates, each containing ~576 µL (96 rxns/plate × 6 µL/rxn) of pooled reactions in 20 mM EDTA.
   a. Note: The most efficient way to do the pooling varies depending on the equipment available. Our group relies on 12-channel multichannel pipets for this step, and so will accomplish pooling by (1) adding 8 µL 100 mM EDTA to each well in a row of a fresh PCR plate, (2) adding 5 µL reaction from each row in the plate-to-be-pooled to this EDTA, and (3) using a single-channel pipet to add 40 µL (leaving 8 µL dead volume) of each of well in the row to a microcentrifuge tube. An alternate strategy might be, for instance, adding 96 µL 100 mM EDTA to a trough, then pipetting 5 µL of all reactions from a plate into this trough. **Whatever strategy is taken, what is important in pooling is that the ratios of the reactions in the pool remain equal—sacrificing some reaction as dead volume is perfectly acceptable to achieve equal mixing in this step.**
9. For each tube made in step 8, take 100 µL of pooled reaction and add it to 20 µL 6x loading dye (NEB B7025S) in a microcentrifuge tube. **It is critical that the loading dye does not contain SDS.** At this point, the remaining pooled reaction from step 8 can be stored at –20 °C for future use (i.e., if the later steps of this protocol ever need to be redone).
   a. Note that most of the pooled reaction is not moved into later steps with this protocol. Again, if relevant automation and molecular biology equipment is available, reactions can be scaled down below 10 µL, reducing wasted reaction. Current reaction sizes are set to minimize pipetting error.
10. Load the contents of each tube made in step 9 into the agarose gel prepared in step 7. The contents of each tube should be kept separate (I.e., loaded into different lanes in the gel). Load a ladder (we typically use 100 bp ladder from NEB, N3231S) in the flanking lanes.
11. Run the agarose gel at 130 V until the bands have sufficiently migrated. Often, you will see two bands: the lower band is usually primer dimer and the upper is the target. Reference the ladder to identify your product, remembering that the two-step PCR adds 120 bp of additional length (from the universal adapter, barcode, and transposase adapters) onto the gene fragment of interest.
12. Gel-extract the target bands from the agarose gel, again keeping bands from different plates separate. We typically use Zymoclean Gel DNA Recovery Kit (Zymo Research, D4001) for this step. Elution should be performed at a low volume—we typically elute in 10 µL of ddH2O.
13. After gel extraction, combine the gel-extracted pools from each plate in equimolar concentrations. We provide a calculator on the evSeq GitHub repository that can be used to normalize *equal-length* fragments to a pre-specified concentration (https://github.com/fhalab/evSeq/tree/master/lib_prep_tools/LibDilCalculator.xlsx).
   a. Note that the quantification here need not be extremely robust. For all results presented in this work, we performed this step using DNA concentrations output by a GE NanoVue Plus.
   b. Tip: It is generally not advised to pool amplicons drastically different in length. Shorter fragments are preferentially sequenced in NGS, and so the shorter amplicon will dominate the number of reads. Separate submissions should be made for libraries with very different lengths.

14. After the previous step, you should have a single tube of cleaned, normalized DNA consisting of all amplicons from all plates to be pooled. This DNA will be submitted to your sequencing provider for inclusion in a multiplexed sequencing run. You should work with your sequencing provider to ensure that all requirements are met to slot into their pipeline. For instance, this protocol assumes that the sequencing provider can add Nextera-compatible Illumina indices and flow-cell-binding sequences via PCR—it should be confirmed that your sequencing provider can do this before submitting your sample.
    a. Note: Throughout this work, we used the "Customized PCR Amplicon Sequencing" services of Laragen Inc. (http://www.laragen.com/laragen_nextgen.php).
    b. Also note that, depending on your sequencing provider, it may be possible (or even necessary) to add the Illumina indices yourself. Again, you should work with your provider to determine the best course of action for submitting evSeq libraries. Adding indices simply requires one final PCR on the pooled evSeq library.
15. Once sequencing is complete, your sequencing provider should return two fastq (or fastq.gz) files to you. One will contain the forward reads for your pooled samples and the other will contain the reverse reads—both files are needed by the evSeq software for processing.
16. Using the files returned in step 15, run the evSeq software to process results and assign variants to their original wells. Detailed instructions on how to use the evSeq software and interpret its outputs are provided on the evSeq Wiki https://fhalab.github.io/evSeq/4-usage.html

### TrpB Site-Saturation Libraries

#### Single-Site-Saturation Library Generation for TrpB

Saturation mutagenesis libraries were prepared using a modification of the "22-codon trick" described by Kille et al.[1] We first designed the following primer templates for each site:

| Site | Direction | Sequence |
|---|---|---|
| 105 | Forward | GGCAAAACCCGTATCATTGCTNNNACGGGTGCTGGTCAGCAC |
| 105 | Reverse | AGCAATGATACGGGTTTTGCCCATTAGTTTTGCCAGCAGAACCTGGC |
| 118 | Forward | GGCGTAGCAACTGCTACCNNNGCAGCGCTGTTCGGTATGGAATGTGTAATCTATATGG |
| 118 | Reverse | GGTAGCAGTTGCTACGCCGTGCTGACCAGC |
| 162 | Forward | GTAAAATCCGGTAGCCGTACCNNNAAAGACGCAATTGACGAAGCTCTG |
| 162 | Reverse | GGTACGGCTACCGGATTTTACCGGTACAACTTTAGCACCCAGCAG |
| 166 | Forward | CGTACCCTGAAAGACGCANNNGACGAAGCTCTGCGTGACTGGATTACCAACC |
| 166 | Reverse | TGCGTCTTTCAGGGTACGGCTACCGGATTTTACCGG |
| 184 | Forward | CTGCAGACCACCTATTACGTGNNNGGCTCTGTGGTTGGTCC |
| 184 | Reverse | CACGTAATAGGTGGTCTGCAGGTTGGTAATCCAGTCACGCAGAGCT |
| 228 | Forward | TACATCGTTGCGTGCGTGNNNGGTGGTTCTAACGCTGCC |
| 228 | Reverse | CACGCACGCAACGATGTAGTCCGGCAGACGGCCTTCT |
| 292 | Forward | GATGACTGGGGTCAAGTTCAGGTGNNNCACTCCGTCTCCGCTG |
| 292 | Reverse | CACCTGAACTTGACCCCAGTCATCCTGCAGAACGAACGTCTTAGAACCG |
| 301 | Forward | TCCGCTGGCCTGGACNNNTCCGGTGTCGGTCCGGA |
| 301 | Reverse | GTCCAGGCCAGCGGAGACGGAGTGGCTCACCTGAACT |

For the forward primers, each sequence of "NNN" was replaced with "NDT", "VHG", and "TGG", resulting in a total of three degenerate primers which could then be mixed at a ratio of 12:9:1, respectively. The reverse primers were used without changes.

We also used primers that bind within the ampicillin resistance (AmpR) gene in pET22b(+) with sequences as follows:

| Site | Direction | Sequence |
|---|---|---|
| AmpR | Forward | CCAACTTACTTCTGACAACGATCGGAGGACCGAAGGAGCTAACCGCTTTTTTGC |
| AmpR | Reverse | CGATCGTTGTCAGAAGTAAGTTGGCCGCAGTGTTATCACTCATGGTTATGGCAG |

These were then paired to the site-specific primers to create two medium-length fragments with a break in the AmpR gene, requiring proper assembly of both fragments together for transformants to grow under ampicillin selection conditions. For the forward site-saturation primers, a PCR was performed using the reverse AmpR primer, resulting in a fragment from ~1500–2000 bp long. For the reverse site-saturation primers, a PCR was performed using the forward AmpR primer, resulting in a fragment ~4500–5000 bp long.

Reactions were prepared as follows:

| Component | Volume/Rxn (µL) |
|---|---|
| $ddH_2O$ | 15.25 |
| Phusion HF Buffer 5x (NEB B0518S) | 5 |
| Mol-Bio Grade DMSO (MP 194819) | 0.5 |
| 10mM dNTPs (NEB N0447) | 0.5 |
| Tm9D8* plasmid (50 ng/µL) | 1 |
| Phusion (NEB M0530) | 0.25 |
| SSM Primer | 1.25 |
| AmpR Primer | 1.25 |

and then subjected to the following thermal cycle, regardless of fragment length:

| Step | Temperature (°C) | Time |
|---|---|---|
| 1 | 98 | 2 min |
| 2 | 65 (Ramp at 10% of max) | 15 s |
| 3 | 72 | 2 min |
| 4 | 98 | 10 s |
| 5 | 65 | 15 s |
| 6 | 72 | 2 min |
| 7 | Return to 4, 30x | |
| 8 | 72 | 5 min |

Once finished, 1 µL of DpnI (NEB R0176S) was added to each of the reactions, which were then incubated at 37 °C for 1 h to digest the unmutated template plasmid. The presence of correctly sized fragments was confirmed via gel electrophoresis and each fragment was then excised from the gel and purified with the Zymoclean Gel DNA Recovery Kit (Zymo Research D4002).

Purified fragments were then assembled following the standard Gibson assembly method.[2] After 1 h at 50 °C, the reaction mixtures were desalted with a DNA Clean & Concentrator-5 kit (Zymo Research D4013) and used to transform electrocompetent E. cloni cells (Lucigen 60051-1). Libraries were spread onto solid agar selection medium consisting of Luria Broth (RPI L24040-5000.0) supplemented with 100 µg/mL carbenicillin ($LB_{carb}$) and incubated at 37 °C until single colonies were observed. Individual colonies were then transferred into the wells of 96-well 2-mL deep-well plates containing 300 µL of $LB_{carb}$ to isolate monoclonal enzyme variants, with 8 wells being reserved for control conditions, giving 4-fold oversampling of the 22-codon library. These cultures were grown overnight at 37 °C, 220 rpm, and 80% humidity in an Infors Multitron HT until they reached stationary phase, at which point 100 µL from each well were mixed with an equal volume of 50% glycerol and stored at –80 °C for future use.

For protein expression, 20 µL of the remaining culture were used to inoculate 630 µL of Terrific Broth with 100 µg/mL carbenicillin ($TB_{carb}$). These were then grown at 37 °C, 220 rpm, and 80% humidity for 3 hours in an Infors Multitron HT, at which point they were placed on ice for 30 minutes. Following this, 50 µL of a

14 mM solution of isopropyl-β-D-thiogalactoside (IPTG; GoldBio #I2481C100) in $TB_{carb}$ were added to each well to induce protein expression at a final concentration of 1 mM IPTG. Expression proceeded in the same Infors Multitron HT shaker as before at 22 °C, 220 rpm for roughly 18 hours. Cells were harvested via centrifugation at 4500*g* for 10 minutes, the supernatant was removed, and the plates (now containing pelleted, expressed cells) were placed at –20 °C until needed.

Once cells had been harvested, cultures for evSeq were prepared. These cultures were started from the 96-well plate glycerol stocks prepared prior to moving into the cell expression protocol; the cultures were grown overnight (~18hrs) in an Infors Multitron HT (220 rpm, 37 °C) to saturation in 96-well deep-well plates in 300 µL of $LB_{carb}$. These cultures were then frozen and stored at –20 °C to be used for sequencing with evSeq.

A GenBank file detailing the plasmid and primers used in this section is available on the evSeq GitHub at https://github.com/fhalab/evSeq/tree/master/genbank_files/tm9d8s.gb.

#### Sequencing TrpB Libraries with evSeq

Frozen overnight cultures were thawed at room temperature. Libraries were then sequenced with the process described above in *evSeq Library Preparation/Data Analysis Protocol*; the evSeq software was run using all default parameters (`average_q_cutoff = 25, bp_q_cutoff = 30, length_cutoff = 0.9, match_score = 1, mismatch_penalty = 0, gap_open_penalty = 3, gap_extension_penalty = 1, variable_thresh = 0.2, variable_count = 10`) with the `return_alignments` flag thrown.

The inner primers used for library preparation are in the table below and sorted based upon the site-saturation positions they were used to sequence.

| Name | Direction | Sites | Sequence |
|---|---|---|---|
| evSeq_102_f | Forward | 105, 118, 162, 166, 184 | CACCCAAGACCACTCTCCGGGCAAAACTAATGGGCAAAACCCG |
| evSeq_184_r | Reverse | 105, 118, 162, 166, 184 | CGGTGTGCGAAGTAGGTGCGATGCGGACCAACCACAGAG |
| evSeq_226_f | Forward | 228, 292, 301 | CACCCAAGACCACTCTCCGGGCCGGACTACATCGTTGCG |
| evSeq_304_r | Reverse | 228, 292, 301 | CGGTGTGCGAAGTAGGTGCCAATAGGCGTGTTCCGGACC |

The barcode plates (Table S3 – Table S10) were paired to positions as follows:

| Position targeted | Barcode plate |
|---|---|
| 105 | DI01 |
| 118 | DI02 |
| 162 | DI03 |
| 166 | DI04 |
| 184 | DI05 |
| 228 | DI06 |
| 292 | DI07 |
| 301 | DI08 |

#### Measuring the Rate of Tryptophan Formation

Rate of tryptophan formation data was collected with the same procedure described in Rix et al. for non-heat-treated lysate preparation in the section “Indole rate measurements” with a few modifications: lysis occurred in 300 µL KPi buffer with 100 µM pyridoxal 5'-phosphate (PLP) supplemented with 1 mg/mL lysozyme, 0.02 mg/mL bovine pancreas DNase I, and 0.1x BugBuster; lysis occurred at 37 °C for 1 h.[3]

### evSeq of *Rma*NOD Combinatorial Libraries

#### Four-Site-Saturation Library Generation for *Rma*NOD

Positions S28, M31, Q52, and L56 of a variant of *Rma*NOD (*Rma*NOD Y32G) were targeted for comprehensive site-saturation mutagenesis using a variant of the 22-codon trick originally described by Kille et al.[1] Due to the proximity of positions S28 and M31, it was easiest to use the same mutagenesis primers to target them; the same was done for positions Q52 and L56. Because the 22-codon trick requires three degenerate codons per position targeted, nine individual primers capturing all combinations (3 codons ^ 2 positions/per primer = 9 primers) of the degenerate codons had to be ordered for each of the two mutagenic primers. Sequences of these primers are given in the table below. Note that the names of the primers are delimited by "-" and that the delimited sections reflect the mutagenized positions, the degenerate codons at those positions, and the direction of the primer on the template DNA ([Positions]-[Codon1]-[Codon2]-[Direction]):

| Name | Sequence |
|---|---|
| S28M31-NDT-NDT-F | AAACACTCAGTCGCTATTNDTGCCACGNDTGGTCGGCTGCTTTTCG |
| S28M31-NDT-VHG-F | AAACACTCAGTCGCTATTNDTGCCACGVHGGGTCGGCTGCTTTTCG |
| S28M31-NDT-TGG-F | AAACACTCAGTCGCTATTNDTGCCACGTGGGGTCGGCTGCTTTTCG |
| S28M31-VHG-NDT-F | AAACACTCAGTCGCTATTVHGGCCACGNDTGGTCGGCTGCTTTTCG |
| S28M31-VHG-VHG-F | AAACACTCAGTCGCTATTVHGGCCACGVHGGGTCGGCTGCTTTTCG |
| S28M31-VHG-TGG-F | AAACACTCAGTCGCTATTVHGGCCACGTGGGGTCGGCTGCTTTTCG |
| S28M31-TGG-NDT-F | AAACACTCAGTCGCTATTTGGGCCACGNDTGGTCGGCTGCTTTTCG |
| S28M31-TGG-VHG-F | AAACACTCAGTCGCTATTTGGGCCACGVHGGGTCGGCTGCTTTTCG |
| S28M31-TGG-TGG-F | AAACACTCAGTCGCTATTTGGGCCACGTGGGGTCGGCTGCTTTTCG |
| Q52L56-AHN-AHN-R | GGCCAACAGGGCCGACGCAHNCTTGTGTATAHNTCTCTCAGGAAGTTCAAACAAG |
| Q52L56-AHN-CDB-R | GGCCAACAGGGCCGACGCAHNCTTGTGTATCDBTCTCTCAGGAAGTTCAAACAAG |
| Q52L56-AHN-CCA-R | GGCCAACAGGGCCGACGCAHNCTTGTGTATCCATCTCTCAGGAAGTTCAAACAAG |
| Q52L56-CDB-AHN-R | GGCCAACAGGGCCGACGCCDBCTTGTGTATAHNTCTCTCAGGAAGTTCAAACAAG |
| Q52L56-CDB-CDB-R | GGCCAACAGGGCCGACGCCDBCTTGTGTATCDBTCTCTCAGGAAGTTCAAACAAG |
| Q52L56-CDB-CCA-R | GGCCAACAGGGCCGACGCCDBCTTGTGTATCCATCTCTCAGGAAGTTCAAACAAG |
| Q52L56-CCA-AHN-R | GGCCAACAGGGCCGACGCCCACTTGTGTATAHNTCTCTCAGGAAGTTCAAACAAG |
| Q52L56-CCA-CDB-R | GGCCAACAGGGCCGACGCCCACTTGTGTATCDBTCTCTCAGGAAGTTCAAACAAG |
| Q52L56-CCA-CCA-R | GGCCAACAGGGCCGACGCCCACTTGTGTATCCATCTCTCAGGAAGTTCAAACAAG |

The above primers were all ordered from IDT at 100 µM. To ensure more equal representation of codons in the final library, the above degenerate codons were combined proportional to the number of individual codons they encoded. Both a "forward" mixture and a "reverse" mixture were prepared according to the below table:

| Primer Mix | Name | Volume (µL) |
|---|---|---|
| Forward | S28M31-NDT-NDT-F | 144 |
| Forward | S28M31-NDT-VHG-F | 108 |
| Forward | S28M31-NDT-TGG-F | 12 |
| Forward | S28M31-VHG-NDT-F | 108 |

| | | |
|---|---|---|
| Forward | S28M31-VHG-VHG-F | 81 |
| Forward | S28M31-VHG-TGG-F | 9 |
| Forward | S28M31-TGG-NDT-F | 12 |
| Forward | S28M31-TGG-VHG-F | 9 |
| Forward | S28M31-TGG-TGG-F | 1 |
| Reverse | Q52L56-AHN-AHN-R | 144 |
| Reverse | Q52L56-AHN-CDB-R | 108 |
| Reverse | Q52L56-AHN-CCA-R | 12 |
| Reverse | Q52L56-CDB-AHN-R | 108 |
| Reverse | Q52L56-CDB-CDB-R | 81 |
| Reverse | Q52L56-CDB-CCA-R | 9 |
| Reverse | Q52L56-CCA-AHN-R | 12 |
| Reverse | Q52L56-CCA-CDB-R | 9 |
| Reverse | Q52L56-CCA-CCA-R | 1 |

A 10 µM forward-reverse primer mixture was prepared by adding 10 µL of both of the above forward and reverse primer mixtures to 80 µL dd$H_2O$. Once the forward-reverse primer mixture was prepared, it was used in a PCR to build a pool of DNA fragments containing the 4-site combinatorial libraries. Two fragments that captured the remainder of the *Rma*NOD gene and host plasmid (pET22b(+)) were also produced by PCR. The primers used for these flanking fragments are given below:

| Flanking Fragment | Primer Type | Primer Name | Sequence |
|---|---|---|---|
| 0 | Forward | Universal-F | CCAACTTACTTCTGACAACGATCGGAGGA CCGAAGGAGCTAACCGCTTTTTTGC |
| 0 | Reverse | S28M31_Const-R | AATAGCGACTGAGTGTTTCTGCAGTGCAG GCAC |
| 1 | Forward | L56_Const-F | GCGTCGGCCCTGTTGGCCTACGCCCGTAG TATCGACAACCC |
| 1 | Reverse | Universal-R | CGATCGTTGTCAGAAGTAAGTTGGCCGCA GTGTTATCACTCATGGTTATGGCAG |

Forward-reverse primer mixtures were also prepared for the flanking primers by adding 10 µL of both forward and reverse primers to 80 µL dd$H_2O$. PCRs for both flanking fragments as well as the variant-containing fragment were prepared as given below:

| Component | Volume / Rxn (µL) |
|---|---|
| Phusion HF Buffer 5x (NEB B0518S) | 10 |
| 10 mM dNTPs (NEB N0447) | 1 |
| Phusion Polymerase (NEB M0530) | 0.5 |
| *Rma*NOD Y32G Plasmid Template (ng/µL) | 0.5 |
| dd$H_2O$ | 35 |
| Mol-Bio Grade DMSO (MP 194819) | 2 |
| Forward-Reverse Primer Mix (10 µM) | 1 |

Cycling times for the all PCRs are given below (note that “TD” means a touchdown PCR):

| Step | Temperature (°C) | Time |
|---|---|---|
| 1 | 98 | 30 s |
| 2 | 98 | 10 s |

| | | |
|---|---|---|
| 3 | 65 - 56 (TD) | 20 s |
| 4 | 72 | 2 min 30 s |
| 5 | Return to 2, 9x | |
| 6 | 98 | 10 s |
| 7 | 65 | 20 s |
| 8 | 72 | 2 min 30 s |
| 9 | Return to 6, 9x | |
| 10 | 72 | 10 min |

After PCR completed 1 μL DpnI (NEB R0176S) was added to each reaction. The reactions were then held at 37 °C in a thermalcycler.

After the DpnI digest, all PCR fragments were gel-extracted using a Zymoclean Gel DNA Recovery Kit (D4002).

Fragments were to eventually be assembled using Gibson assembly.[2] Because the efficiency of Gibson assembly increases with decreasing numbers of fragments, an assembly PCR was performed to combine flanking fragment 1 and the variant fragment. To begin, a 5 ng/μL template mix of gel-extracted flanking fragment 1 and the variant fragment was prepared; a 10 μM primer mix was also prepared by adding 10 μL of both the 100 μM forward primer for flanking fragment 1 and 10 μL of the 100 μM forward pool of primers prepared for the variant fragment. The assembly PCR was then set up as below:

| **Component** | **Volume (uL)** |
|---|---|
| Phusion HF Buffer 5x (NEB B0518S) | 10 |
| 10 mM dNTPs (NEB N0447) | 1 |
| Assembly 10 μM Primer Mixture | 1 |
| Phusion Polymerase (NEB M0530) | 0.5 |
| Assembly 5 ng/μL PCR Product Template Mixture | 0.5 |
| dd$H_2O$ | 35 |
| Mol-Bio Grade DMSO (MP 194819) | 2 |

The cycling times for the assembly PCR is given below:

| **Step** | **Temperature (°C)** | **Time** |
|---|---|---|
| 1 | 98 | 30 s |
| 2 | 98 | 10 s |
| 3 | 60 - 51 (TD) | 20 s |
| 4 | 72 | 1 min |
| 5 | Return to 2, 9x | |
| 6 | 98 | 10 s |
| 7 | 65 | 20 s |
| 8 | 72 | 1 min |
| 9 | Return to 6, 14x | |
| 10 | 72 | 10 min |

The resultant assembled fragment was then gel-extracted, again using a Zymoclean Gel DNA Recovery Kit (D4002).

To complete construction of the library of variant plasmids, a Gibson assembly was performed to combine the assembled PCR fragment and flanking fragment 0.[2] After Gibson assembly, the Gibson reaction was cleaned using a Monarch PCR & DNA Cleanup Kit (NEB CAT T1030L).

The cleaned Gibson product was next used to transform electrocompetent E. cloni BL21 DE3. In detail, a cuvette with a 2 mm path length (USA CAT 9104-5050) was placed on ice to cool. Next, 150 μL of electrocompetent BL21 DE3 were thawed on ice. Then, 6 μL of cleaned Gibson product were added to the thawed aliquot; after addition of Gibson product, the aliquot was immediately moved into the cooled cuvette.

The cells were electroporated with a Bio-Rad GenePulser XCell at 2.5 kV before being immediately resuspended in in 750 µL LB. Immediately after resuspension (i.e., with no recovery), cells were diluted 3-fold in LB + ampicillin before 100 µL and 50 µL resuspension was plated on 2 separate LB + 100 µg/mL ampicillin agar plates. The agar plates were left in a 37 °C incubator for 14 h.

To build the 96-well plates of *Rma*NOD variants used to demonstrate evSeq, 400 µL LB + 100 µg/mL ampicillin was first added to each well of 5x 96-well deepwell plates. Colonies from the agar plates grown overnight were then picked into the wells of the deepwell plates. The plates were sealed with breathable film before being placed in an Infors Multitron HT at 240 rpm, 37 °C for ~16 h. To glycerol stock the now-stationary-phase culture, 100 µL overnight culture was added to 100 µL 50% glycerol before being stored at -80 °C until its use in evSeq library preparation.

A GenBank file detailing the plasmid and primers used in this section is available on the evSeq GitHub (https://github.com/fhalab/evSeq/tree/master/genbank_files/rmanod_y32g.gb).

#### Sequencing *Rma*NOD libraries with evSeq

To begin preparation of culture for evSeq with the *Rma*NOD variants, cultures in 96-well deep-well plates (with 300 µL of $LB_{carb}$) were started from the 96-well plate glycerol stocks prepared in the previous section. The plates were sealed with breathable film before being placed in an Infors Multitron HT at 240 rpm; the cultures were grown overnight (~18hrs) before being frozen and stored at –20 °C.

To start the evSeq protocol, frozen overnight cultures were thawed in a room temperature water bath. Libraries were then sequenced with the process described in the section *evSeq Library Preparation/Data Analysis Protocol*, above; the evSeq software was run using the same parameters as for the TrpB data analysis (see *Sequencing TrpB Libraries with evSeq*, above).

The inner primers used for evSeq of the *Rma*NOD libraries are in the table below:

| Plates | Forward primer | Reverse Primer |
|---|---|---|
| All plates | CACCCAAGACCACTCTCCGGCACTGCAGAAACACTCAGTCG | CGGTGTGCGAAGTAGGTGCACTACGGGCGTAGGCCAAC |

The barcode plates were paired to plates of variants as follows.

| Position targeted | Barcode plate |
|---|---|
| Plate #1 | DI01 |
| Plate #2 | DI02 |
| Plate #3 | DI03 |
| Plate #4 | DI04 |
| Plate #5 | DI05 |

### Supplemental Figures

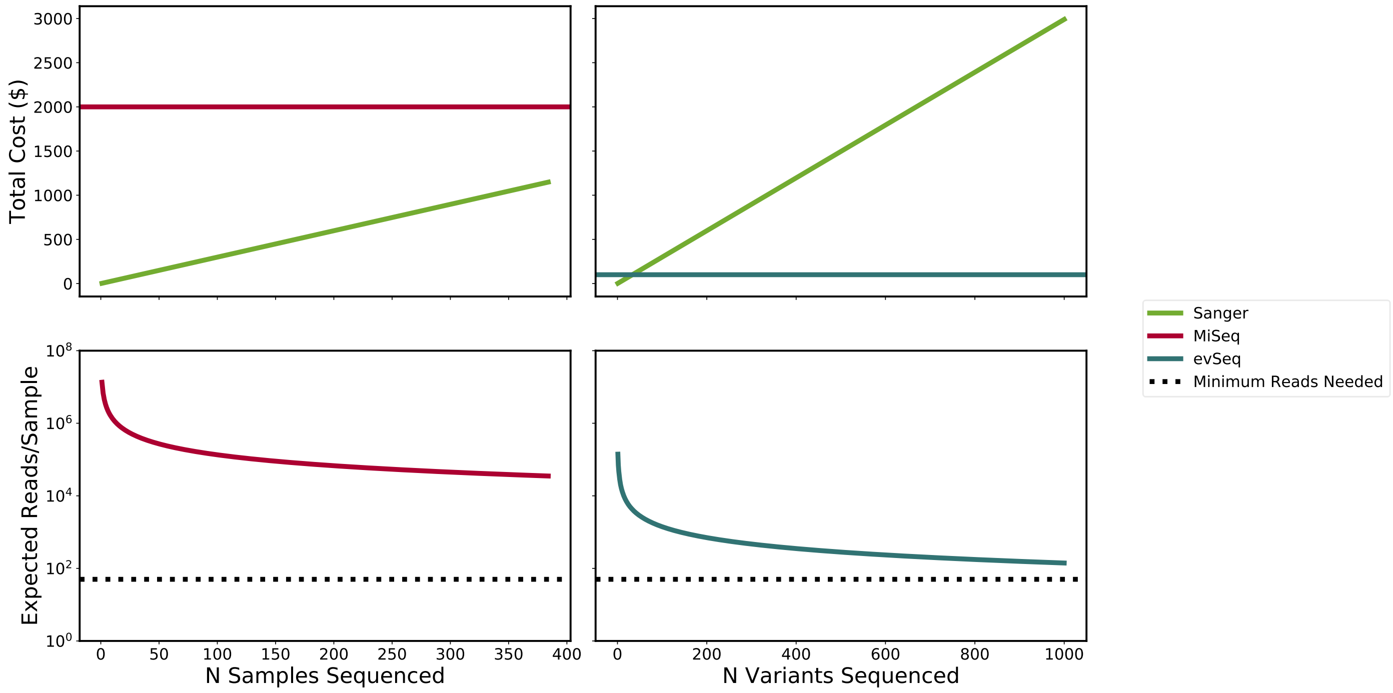

**Figure S1.** Comparison of the tradeoff between sequencing depth and cost for Sanger sequencing (green), a multiplexed MiSeq run (red), and an evSeq library (blue). The top row gives the total cost for sequencing a given number of variants; the bottom row gives the expected number of reads per variant for sequencing a given number of variants. Note that the x-axes for the left and right columns are different. The limit on the x-axis for the left column is set to reflect what is typically the maximum level of multiplexed NGS available (384 samples) when outsourcing sequencing. To be consistent with the language used throughout the main text, the x-axis labels refer to elements run in a multiplexed NGS run as “samples” and elements contained in an evSeq library as “variants”. We assume that the elements sequenced in these examples are derived from protein mutant libraries amenable to sequencing by evSeq (i.e., the sequenced elements are targeted amplicons). **Top Row**: We see that both multiplexed NGS on a commercial MiSeq run and evSeq have constant cost with an increasing number of elements sequenced; Sanger, in contrast, scales linearly with the number of elements sequenced. Many elements (669 with the cost estimates used to make this figure) need to be added to a multiplexed MiSeq run before it becomes more cost-effective than Sanger. Even though research groups may frequently meet or exceed 669 variants in a standard protein engineering experiment, the flat cost of $2000 is far too high to justify regular sequencing of every variant. Many fewer variants (34) need to be added to an evSeq run before it becomes cost-effective over Sanger. A flat cost of ~$100 is justifiable for regularly sequencing all variants. **Bottom Row:** NGS technologies trade off sequencing depth for cost effectiveness. Notably, the per-sample sequencing depth achieved by commercially available multiplexed runs is much higher than what is needed for reliable sequencing. evSeq, in contrast, more efficiently spreads reads, keeping the expected number of reads closer to, yet still above the minimum needed for effective sequencing. **Notes on Figure Generation:** Cost of a single MiSeq run ($2000) is based on an estimate provided by Laragen Inc. Cost of a single Sanger sequencing run ($2.99) is based on a quote from MCLAB for sequencing a single 96-well plate. The number of expected reads from a MiSeq run (13.5 million) is based on estimates provided by Illumina for a MiSeq Reagent Kit v2 (note that almost double the number of reads can be achieved using a v3 kit—we used v2 here to be conservative with our estimates for NGS/evSeq). The number of expected reads for a variant sampled with evSeq assumes the evSeq library was sequenced as 1 of 96 samples on a multiplexed sequencing run using a MiSeq Reagent Kit v2. The cost of a single evSeq run is based on an estimate provided by Laragen for a single sample in a multiplexed sequencing run using a PE150 kit.

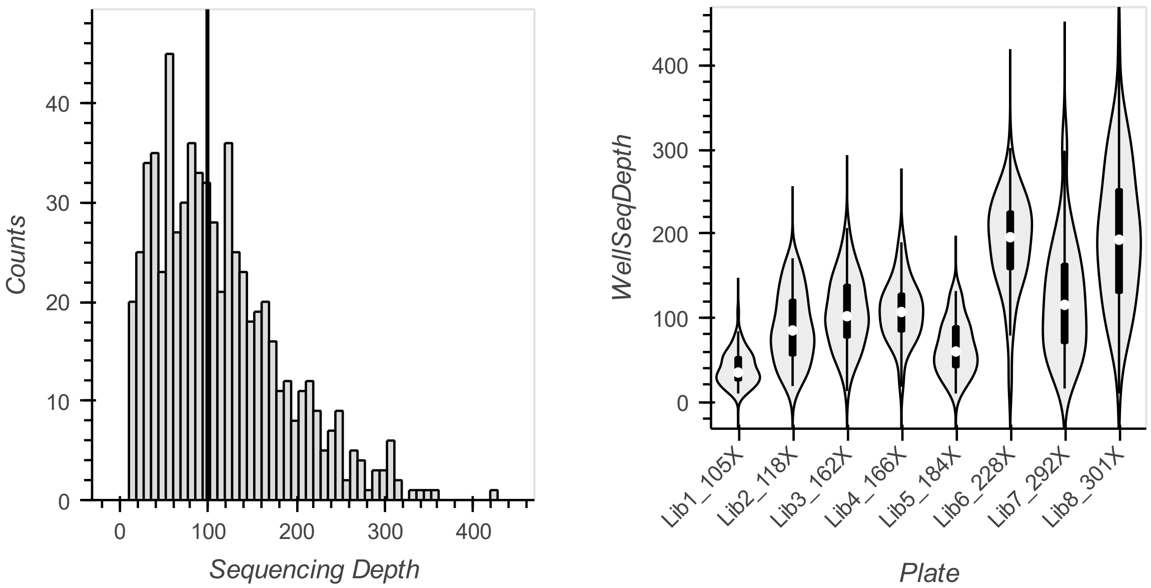

**Figure S2.** Sequencing depths for the *Tm*TrpB9D8* evSeq libraries. **Left:** A histogram of sequencing depths for each *Tm*TrpB9D8* variant contained in the full evSeq library. The vertical black line gives the median. **Right:** Violin plots showing the distribution of read depths over the wells in each sequenced plate. Variability between plates likely indicates inaccurate quantification of pooled plates prior to final assembly of the evSeq library. Notable, libraries 1-5 use different evSeq primers than libraries 6-8.

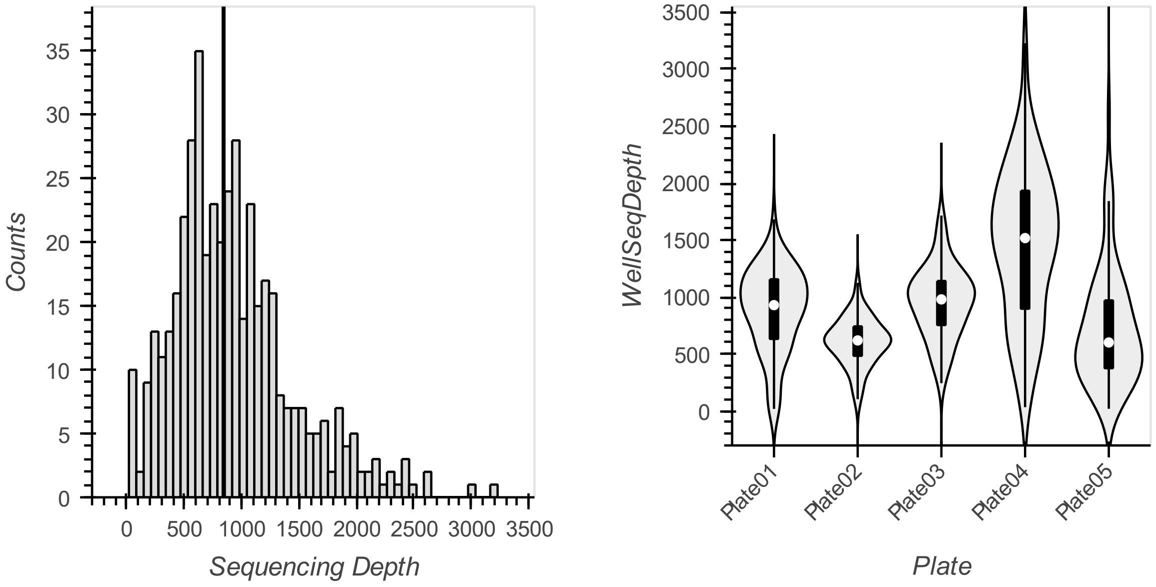

**Figure S3.** Sequencing depths for the *Rma*NOD evSeq libraries. **Left:** A histogram of sequencing depths for each *Rma*NOD variant contained in the full evSeq library. The vertical black line gives the median. **Right:** Violin plots showing the distribution of read depths over the wells in each sequenced plate. Variability between plates likely indicates inaccurate quantification of pooled plates prior to final assembly of the evSeq library.

#### Barcode and Outer Primer Sequences

**Table S1.** evSeq barcode sequences used in this work. The "Plate" and "Well" columns give the location of these sequences in the IDT order form provided on the evSeq GitHub repository (see *Ordering Barcode Primers from IDT* and *Barcode Design*, above). Note that barcode sequences can also be found in the "index_map.csv" file found on the evSeq GitHub repository (https://github.com/fhalab/evSeq/tree/master/evSeq/util/index_map.csv); this csv file also gives the combinations of barcodes used to define the dual indexing (DI) plates.

| Plate | Well | Barcode |
|---|---|---|
| FBC | A01 | GATCATG |
| FBC | A02 | TACATGG |
| FBC | A03 | AAGCACC |
| FBC | A04 | TGGCTCA |
| FBC | A05 | CTTGCTC |
| FBC | A06 | GAAGCGT |
| FBC | A07 | TCTCCAT |
| FBC | A08 | TTGAAGG |
| FBC | A09 | GAATGTC |
| FBC | A10 | ATCTCCA |
| FBC | A11 | GCGTTAT |
| FBC | A12 | TGCACCA |
| FBC | B01 | TGCCTAT |
| FBC | B02 | AGGAATC |
| FBC | B03 | TCCACTG |
| FBC | B04 | TTGTACC |
| FBC | B05 | TTCGAGT |
| FBC | B06 | CTTCAGC |
| FBC | B07 | CAGTGCA |
| FBC | B08 | TGCTGTC |
| FBC | B09 | CGCCATT |
| FBC | B10 | GCCATGA |
| FBC | B11 | CACAACG |
| FBC | B12 | CTTCGCT |
| FBC | C01 | TCGTGAA |
| FBC | C02 | TTATCGG |
| FBC | C03 | AGACCAT |
| FBC | C04 | ACATGAG |
| FBC | C05 | ACGTACT |
| FBC | C06 | CACCTCA |
| FBC | C07 | GTTGGAG |
| FBC | C08 | TGTTCTG |
| FBC | C09 | CTTACGT |
| FBC | C10 | GAGGTTG |
| FBC | C11 | ATGGACA |
| FBC | C12 | ACACTGA |

| | | |
|---|---|---|
| FBC | D01 | ATCTGTG |
| FBC | D02 | AATGTGC |
| FBC | D03 | GAGTTGA |
| FBC | D04 | TTCTCAC |
| FBC | D05 | TGAAGCG |
| FBC | D06 | GCTACAA |
| FBC | D07 | AGAGAAC |
| FBC | D08 | CAGAGTG |
| FBC | D09 | TTCCGAA |
| FBC | D10 | GTACGAC |
| FBC | D11 | ACTCTTG |
| FBC | D12 | CCAACCA |
| FBC | E01 | CTCTAGA |
| FBC | E02 | AATCGGA |
| FBC | E03 | CGTCCTA |
| FBC | E04 | GGAATGT |
| FBC | E05 | TCCAAGC |
| FBC | E06 | GCACCTA |
| FBC | E07 | TTGCGTT |
| FBC | E08 | CAGGATT |
| FBC | E09 | CTGCATA |
| FBC | E10 | CGTTGAG |
| FBC | E11 | TGCTACT |
| FBC | E12 | GTGATCC |
| FBC | F01 | GCATGGT |
| FBC | F02 | GTCGTTA |
| FBC | F03 | CCTGACA |
| FBC | F04 | AGTGTAG |
| FBC | F05 | CGAGCAA |
| FBC | F06 | CTACTCC |
| FBC | F07 | GATGCCA |
| FBC | F08 | GACCGAT |
| FBC | F09 | ACGTTGG |
| FBC | F10 | ATGAGCG |
| FBC | F11 | TACTCCG |
| FBC | F12 | GATTCAC |
| FBC | G01 | ATGACGC |
| FBC | G02 | GGTTGTT |
| FBC | G03 | GTACTTG |
| FBC | G04 | TAGCAAG |
| FBC | G05 | CTGCCAT |
| FBC | G06 | GAGAACA |
| FBC | G07 | GTATAGC |

| | | |
|---|---|---|
| FBC | G08 | TGATGGA |
| FBC | G09 | GGCAGTA |
| FBC | G10 | GAAGAAG |
| FBC | G11 | AGCGGTT |
| FBC | G12 | TAAGGCC |
| FBC | H01 | AACCTGT |
| FBC | H02 | AGTACAC |
| FBC | H03 | CTCGTAG |
| FBC | H04 | CTAGGTG |
| FBC | H05 | CGATACC |
| FBC | H06 | TCGGCTA |
| FBC | H07 | CGGTTGT |
| FBC | H08 | ATTGCCT |
| FBC | H09 | CATTCGA |
| FBC | H10 | GCACAAT |
| FBC | H11 | GCAGTAA |
| FBC | H12 | CCTAATC |
| RBC | A01 | GAACTGC |
| RBC | A02 | ACCAGGT |
| RBC | A03 | TCTAGAG |
| RBC | A04 | CACACAA |
| RBC | A05 | GTGGAAC |
| RBC | A06 | ATATGCC |
| RBC | A07 | GGTCTGA |
| RBC | A08 | GTGAGAT |
| RBC | A09 | TTGGCAG |
| RBC | A10 | ATGCCTG |
| RBC | A11 | TCCGAAG |
| RBC | A12 | GGCTTAC |
| RBC | B01 | AGTTGGC |
| RBC | B02 | AACGATG |
| RBC | B03 | ACTACCG |
| RBC | B04 | GGTGTCT |
| RBC | B05 | CCAGCTT |
| RBC | B06 | TTAGACG |
| RBC | B07 | ACCATAC |
| RBC | B08 | GACGACT |
| RBC | B09 | GTCACCT |
| RBC | B10 | CGTGATG |
| RBC | B11 | GCTTCCT |
| RBC | B12 | TAGACGT |
| RBC | C01 | CGGACTT |
| RBC | C02 | ACCGGAA |

| | | |
|---|---|---|
| RBC | C03 | CCGAAGT |
| RBC | C04 | TCACGCA |
| RBC | C05 | ATCCTCG |
| RBC | C06 | CGAATAG |
| RBC | C07 | TATCCGG |
| RBC | C08 | AGCAAGA |
| RBC | C09 | TGTCGAC |
| RBC | C10 | TTCCATG |
| RBC | C11 | GCAATCG |
| RBC | C12 | TGAGTGG |
| RBC | D01 | TAGGAGA |
| RBC | D02 | AGTCAGT |
| RBC | D03 | GTGCTGT |
| RBC | D04 | CAACAAC |
| RBC | D05 | AATAGCC |
| RBC | D06 | TCTGTGA |
| RBC | D07 | TGTGGTA |
| RBC | D08 | GCGTATG |
| RBC | D09 | AGTTACG |
| RBC | D10 | TTCCTGC |
| RBC | D11 | TATGTCG |
| RBC | D12 | GGAGAGA |
| RBC | E01 | CCTTAGG |
| RBC | E02 | TGTATCC |
| RBC | E03 | CAACCTG |
| RBC | E04 | CTGATGA |
| RBC | E05 | AAGACAG |
| RBC | E06 | AGCTCGT |
| RBC | E07 | GATTGCG |
| RBC | E08 | TCCTTCA |
| RBC | E09 | TCACAGG |
| RBC | E10 | AGAGCTG |
| RBC | E11 | CCTCTGT |
| RBC | E12 | CCTCGAA |
| RBC | F01 | GTGTCTC |
| RBC | F02 | ATTGAGG |
| RBC | F03 | GACAATC |
| RBC | F04 | CACTTGC |
| RBC | F05 | TGAACGC |
| RBC | F06 | CGTAGCA |
| RBC | F07 | AGGTTCC |
| RBC | F08 | GTACACA |
| RBC | F09 | GATAGGT |

| | | |
|---|---|---|
| RBC | F10 | TAGCCTC |
| RBC | F11 | TTCAGCC |
| RBC | F12 | GGATTCA |
| RBC | G01 | TGAGCCT |
| RBC | G02 | AACGCGA |
| RBC | G03 | TCATTGC |
| RBC | G04 | AGCATCT |
| RBC | G05 | TTGGTCT |
| RBC | G06 | CAAGGAT |
| RBC | G07 | AGACGTC |
| RBC | G08 | AGGTCAA |
| RBC | G09 | ATGCTAC |
| RBC | G10 | CTCTGAT |
| RBC | G11 | TCAAGTC |
| RBC | G12 | TCGAGCT |
| RBC | H01 | ACAGTCT |
| RBC | H02 | CAGATAC |
| RBC | H03 | TACGTTC |
| RBC | H04 | ACGGTTC |
| RBC | H05 | CATCGTC |
| RBC | H06 | TACGCAT |
| RBC | H07 | CTTAGAC |
| RBC | H08 | AACTGAC |
| RBC | H09 | ACTTGCA |
| RBC | H10 | ACGCGAT |
| RBC | H11 | TCGACAC |
| RBC | H12 | ACTCAAC |

**Table S2.** Full-length evSeq barcode (outer) primer sequences used in this work. The "Plate" and "Well" columns give the location of these sequences in the IDT order form provided on the evSeq GitHub repository (see *Ordering Barcode Primers from IDT* and *Preparation of evSeq Barcode Primer Mixes*, above).

| Plate | Well | Sequence |
|---|---|---|
| FBC | A01 | TCGTCGGCAGCGTCAGATGTGTATAAGAGACAGGATCATGCACCCAAGACCACTCTCCGG |
| FBC | A02 | TCGTCGGCAGCGTCAGATGTGTATAAGAGACAGTACATGGCACCCAAGACCACTCTCCGG |
| FBC | A03 | TCGTCGGCAGCGTCAGATGTGTATAAGAGACAGAAGCACCCACCCAAGACCACTCTCCGG |
| FBC | A04 | TCGTCGGCAGCGTCAGATGTGTATAAGAGACAGTGGCTCACACCCAAGACCACTCTCCGG |
| FBC | A05 | TCGTCGGCAGCGTCAGATGTGTATAAGAGACAGCTTGCTCCACCCAAGACCACTCTCCGG |
| FBC | A06 | TCGTCGGCAGCGTCAGATGTGTATAAGAGACAGGAAGCGTCACCCAAGACCACTCTCCGG |
| FBC | A07 | TCGTCGGCAGCGTCAGATGTGTATAAGAGACAGTCTCCATCACCCAAGACCACTCTCCGG |
| FBC | A08 | TCGTCGGCAGCGTCAGATGTGTATAAGAGACAGTTGAAGGCACCCAAGACCACTCTCCGG |
| FBC | A09 | TCGTCGGCAGCGTCAGATGTGTATAAGAGACAGGAATGTCCACCCAAGACCACTCTCCGG |
| FBC | A10 | TCGTCGGCAGCGTCAGATGTGTATAAGAGACAGATCTCCACACCCAAGACCACTCTCCGG |
| FBC | A11 | TCGTCGGCAGCGTCAGATGTGTATAAGAGACAGGCGTTATCACCCAAGACCACTCTCCGG |

| | | |
|---|---|---|
| FBC | A12 | TCGTCGGCAGCGTCAGATGTGTATAAGAGACAGTGCACCACACCCAAGACCACTCTCCGG |
| FBC | B01 | TCGTCGGCAGCGTCAGATGTGTATAAGAGACAGTGCCTATCACCCAAGACCACTCTCCGG |
| FBC | B02 | TCGTCGGCAGCGTCAGATGTGTATAAGAGACAGAGGAATCCACCCAAGACCACTCTCCGG |
| FBC | B03 | TCGTCGGCAGCGTCAGATGTGTATAAGAGACAGTCCACTGCACCCAAGACCACTCTCCGG |
| FBC | B04 | TCGTCGGCAGCGTCAGATGTGTATAAGAGACAGTTGTACCCACCCAAGACCACTCTCCGG |
| FBC | B05 | TCGTCGGCAGCGTCAGATGTGTATAAGAGACAGTTCGAGTCACCCAAGACCACTCTCCGG |
| FBC | B06 | TCGTCGGCAGCGTCAGATGTGTATAAGAGACAGCTTCAGCCACCCAAGACCACTCTCCGG |
| FBC | B07 | TCGTCGGCAGCGTCAGATGTGTATAAGAGACAGCAGTGCACACCCAAGACCACTCTCCGG |
| FBC | B08 | TCGTCGGCAGCGTCAGATGTGTATAAGAGACAGTGCTGTCCACCCAAGACCACTCTCCGG |
| FBC | B09 | TCGTCGGCAGCGTCAGATGTGTATAAGAGACAGCGCCATTCACCCAAGACCACTCTCCGG |
| FBC | B10 | TCGTCGGCAGCGTCAGATGTGTATAAGAGACAGGCCATGACACCCAAGACCACTCTCCGG |
| FBC | B11 | TCGTCGGCAGCGTCAGATGTGTATAAGAGACAGCACAACGCACCCAAGACCACTCTCCGG |
| FBC | B12 | TCGTCGGCAGCGTCAGATGTGTATAAGAGACAGCTTCGCTCACCCAAGACCACTCTCCGG |
| FBC | C01 | TCGTCGGCAGCGTCAGATGTGTATAAGAGACAGTCGTGAACACCCAAGACCACTCTCCGG |
| FBC | C02 | TCGTCGGCAGCGTCAGATGTGTATAAGAGACAGTTATCGGCACCCAAGACCACTCTCCGG |
| FBC | C03 | TCGTCGGCAGCGTCAGATGTGTATAAGAGACAGAGACCATCACCCAAGACCACTCTCCGG |
| FBC | C04 | TCGTCGGCAGCGTCAGATGTGTATAAGAGACAGACATGAGCACCCAAGACCACTCTCCGG |
| FBC | C05 | TCGTCGGCAGCGTCAGATGTGTATAAGAGACAGACGTACTCACCCAAGACCACTCTCCGG |
| FBC | C06 | TCGTCGGCAGCGTCAGATGTGTATAAGAGACAGCACCTCACACCCAAGACCACTCTCCGG |
| FBC | C07 | TCGTCGGCAGCGTCAGATGTGTATAAGAGACAGGTTGGAGCACCCAAGACCACTCTCCGG |
| FBC | C08 | TCGTCGGCAGCGTCAGATGTGTATAAGAGACAGTGTTCTGCACCCAAGACCACTCTCCGG |
| FBC | C09 | TCGTCGGCAGCGTCAGATGTGTATAAGAGACAGCTTACGTCACCCAAGACCACTCTCCGG |
| FBC | C10 | TCGTCGGCAGCGTCAGATGTGTATAAGAGACAGGAGGTTGCACCCAAGACCACTCTCCGG |
| FBC | C11 | TCGTCGGCAGCGTCAGATGTGTATAAGAGACAGATGGACACACCCAAGACCACTCTCCGG |
| FBC | C12 | TCGTCGGCAGCGTCAGATGTGTATAAGAGACAGACACTGACACCCAAGACCACTCTCCGG |
| FBC | D01 | TCGTCGGCAGCGTCAGATGTGTATAAGAGACAGATCTGTGCACCCAAGACCACTCTCCGG |
| FBC | D02 | TCGTCGGCAGCGTCAGATGTGTATAAGAGACAGAATGTGCCACCCAAGACCACTCTCCGG |
| FBC | D03 | TCGTCGGCAGCGTCAGATGTGTATAAGAGACAGGAGTTGACACCCAAGACCACTCTCCGG |
| FBC | D04 | TCGTCGGCAGCGTCAGATGTGTATAAGAGACAGTTCTCACCACCCAAGACCACTCTCCGG |
| FBC | D05 | TCGTCGGCAGCGTCAGATGTGTATAAGAGACAGTGAAGCGCACCCAAGACCACTCTCCGG |
| FBC | D06 | TCGTCGGCAGCGTCAGATGTGTATAAGAGACAGGCTACAACACCCAAGACCACTCTCCGG |
| FBC | D07 | TCGTCGGCAGCGTCAGATGTGTATAAGAGACAGAGAGAACCACCCAAGACCACTCTCCGG |
| FBC | D08 | TCGTCGGCAGCGTCAGATGTGTATAAGAGACAGCAGAGTGCACCCAAGACCACTCTCCGG |
| FBC | D09 | TCGTCGGCAGCGTCAGATGTGTATAAGAGACAGTTCCGAACACCCAAGACCACTCTCCGG |
| FBC | D10 | TCGTCGGCAGCGTCAGATGTGTATAAGAGACAGGTACGACCACCCAAGACCACTCTCCGG |
| FBC | D11 | TCGTCGGCAGCGTCAGATGTGTATAAGAGACAGACTCTTGCACCCAAGACCACTCTCCGG |
| FBC | D12 | TCGTCGGCAGCGTCAGATGTGTATAAGAGACAGCCAACCACACCCAAGACCACTCTCCGG |
| FBC | E01 | TCGTCGGCAGCGTCAGATGTGTATAAGAGACAGCTCTAGACACCCAAGACCACTCTCCGG |
| FBC | E02 | TCGTCGGCAGCGTCAGATGTGTATAAGAGACAGAATCGGACACCCAAGACCACTCTCCGG |
| FBC | E03 | TCGTCGGCAGCGTCAGATGTGTATAAGAGACAGCGTCCTACACCCAAGACCACTCTCCGG |
| FBC | E04 | TCGTCGGCAGCGTCAGATGTGTATAAGAGACAGGGAATGTCACCCAAGACCACTCTCCGG |
| FBC | E05 | TCGTCGGCAGCGTCAGATGTGTATAAGAGACAGTCCAAGCCACCCAAGACCACTCTCCGG |
| FBC | E06 | TCGTCGGCAGCGTCAGATGTGTATAAGAGACAGGCACCTACACCCAAGACCACTCTCCGG |

| | | |
|---|---|---|
| FBC | E07 | TCGTCGGCAGCGTCAGATGTGTATAAGAGACAGTTGCGTTCACCCAAGACCACTCTCCGG |
| FBC | E08 | TCGTCGGCAGCGTCAGATGTGTATAAGAGACAGCAGGATTCACCCAAGACCACTCTCCGG |
| FBC | E09 | TCGTCGGCAGCGTCAGATGTGTATAAGAGACAGCTGCATACACCCAAGACCACTCTCCGG |
| FBC | E10 | TCGTCGGCAGCGTCAGATGTGTATAAGAGACAGCGTTGAGCACCCAAGACCACTCTCCGG |
| FBC | E11 | TCGTCGGCAGCGTCAGATGTGTATAAGAGACAGTGCTACTCACCCAAGACCACTCTCCGG |
| FBC | E12 | TCGTCGGCAGCGTCAGATGTGTATAAGAGACAGGTGATCCCACCCAAGACCACTCTCCGG |
| FBC | F01 | TCGTCGGCAGCGTCAGATGTGTATAAGAGACAGGCATGGTCACCCAAGACCACTCTCCGG |
| FBC | F02 | TCGTCGGCAGCGTCAGATGTGTATAAGAGACAGGTCGTTACACCCAAGACCACTCTCCGG |
| FBC | F03 | TCGTCGGCAGCGTCAGATGTGTATAAGAGACAGCCTGACACACCCAAGACCACTCTCCGG |
| FBC | F04 | TCGTCGGCAGCGTCAGATGTGTATAAGAGACAGAGTGTAGCACCCAAGACCACTCTCCGG |
| FBC | F05 | TCGTCGGCAGCGTCAGATGTGTATAAGAGACAGCGAGCAACACCCAAGACCACTCTCCGG |
| FBC | F06 | TCGTCGGCAGCGTCAGATGTGTATAAGAGACAGCTACTCCCACCCAAGACCACTCTCCGG |
| FBC | F07 | TCGTCGGCAGCGTCAGATGTGTATAAGAGACAGGATGCCACACCCAAGACCACTCTCCGG |
| FBC | F08 | TCGTCGGCAGCGTCAGATGTGTATAAGAGACAGGACCGATCACCCAAGACCACTCTCCGG |
| FBC | F09 | TCGTCGGCAGCGTCAGATGTGTATAAGAGACAGACGTTGGCACCCAAGACCACTCTCCGG |
| FBC | F10 | TCGTCGGCAGCGTCAGATGTGTATAAGAGACAGATGAGCGCACCCAAGACCACTCTCCGG |
| FBC | F11 | TCGTCGGCAGCGTCAGATGTGTATAAGAGACAGTACTCCGCACCCAAGACCACTCTCCGG |
| FBC | F12 | TCGTCGGCAGCGTCAGATGTGTATAAGAGACAGGATTCACCACCCAAGACCACTCTCCGG |
| FBC | G01 | TCGTCGGCAGCGTCAGATGTGTATAAGAGACAGATGACGCCACCCAAGACCACTCTCCGG |
| FBC | G02 | TCGTCGGCAGCGTCAGATGTGTATAAGAGACAGGGTTGTTCACCCAAGACCACTCTCCGG |
| FBC | G03 | TCGTCGGCAGCGTCAGATGTGTATAAGAGACAGGTACTTGCACCCAAGACCACTCTCCGG |
| FBC | G04 | TCGTCGGCAGCGTCAGATGTGTATAAGAGACAGTAGCAAGCACCCAAGACCACTCTCCGG |
| FBC | G05 | TCGTCGGCAGCGTCAGATGTGTATAAGAGACAGCTGCCATCACCCAAGACCACTCTCCGG |
| FBC | G06 | TCGTCGGCAGCGTCAGATGTGTATAAGAGACAGGAGAACACACCCAAGACCACTCTCCGG |
| FBC | G07 | TCGTCGGCAGCGTCAGATGTGTATAAGAGACAGGTATAGCCACCCAAGACCACTCTCCGG |
| FBC | G08 | TCGTCGGCAGCGTCAGATGTGTATAAGAGACAGTGATGGACACCCAAGACCACTCTCCGG |
| FBC | G09 | TCGTCGGCAGCGTCAGATGTGTATAAGAGACAGGGCAGTACACCCAAGACCACTCTCCGG |
| FBC | G10 | TCGTCGGCAGCGTCAGATGTGTATAAGAGACAGGAAGAAGCACCCAAGACCACTCTCCGG |
| FBC | G11 | TCGTCGGCAGCGTCAGATGTGTATAAGAGACAGAGCGGTTCACCCAAGACCACTCTCCGG |
| FBC | G12 | TCGTCGGCAGCGTCAGATGTGTATAAGAGACAGTAAGGCCCACCCAAGACCACTCTCCGG |
| FBC | H01 | TCGTCGGCAGCGTCAGATGTGTATAAGAGACAGAACCTGTCACCCAAGACCACTCTCCGG |
| FBC | H02 | TCGTCGGCAGCGTCAGATGTGTATAAGAGACAGAGTACACCACCCAAGACCACTCTCCGG |
| FBC | H03 | TCGTCGGCAGCGTCAGATGTGTATAAGAGACAGCTCGTAGCACCCAAGACCACTCTCCGG |
| FBC | H04 | TCGTCGGCAGCGTCAGATGTGTATAAGAGACAGCTAGGTGCACCCAAGACCACTCTCCGG |
| FBC | H05 | TCGTCGGCAGCGTCAGATGTGTATAAGAGACAGCGATACCCACCCAAGACCACTCTCCGG |
| FBC | H06 | TCGTCGGCAGCGTCAGATGTGTATAAGAGACAGTCGGCTACACCCAAGACCACTCTCCGG |
| FBC | H07 | TCGTCGGCAGCGTCAGATGTGTATAAGAGACAGCGGTTGTCACCCAAGACCACTCTCCGG |
| FBC | H08 | TCGTCGGCAGCGTCAGATGTGTATAAGAGACAGATTGCCTCACCCAAGACCACTCTCCGG |
| FBC | H09 | TCGTCGGCAGCGTCAGATGTGTATAAGAGACAGCATTCGACACCCAAGACCACTCTCCGG |
| FBC | H10 | TCGTCGGCAGCGTCAGATGTGTATAAGAGACAGGCACAATCACCCAAGACCACTCTCCGG |
| FBC | H11 | TCGTCGGCAGCGTCAGATGTGTATAAGAGACAGGCAGTAACACCCAAGACCACTCTCCGG |
| FBC | H12 | TCGTCGGCAGCGTCAGATGTGTATAAGAGACAGCCTAATCCACCCAAGACCACTCTCCGG |
| RBC | A01 | GTCTCGTGGGCTCGGAGATGTGTATAAGAGACAGGAACTGCCGGTGTGCGAAGTAGGTGC |

| | | |
|---|---|---|
| RBC | A02 | GTCTCGTGGGCTCGGAGATGTGTATAAGAGACAGACCAGGTCGGTGTGCGAAGTAGGTGC |
| RBC | A03 | GTCTCGTGGGCTCGGAGATGTGTATAAGAGACAGTCTAGAGCGGTGTGCGAAGTAGGTGC |
| RBC | A04 | GTCTCGTGGGCTCGGAGATGTGTATAAGAGACAGCACACAACGGTGTGCGAAGTAGGTGC |
| RBC | A05 | GTCTCGTGGGCTCGGAGATGTGTATAAGAGACAGGTGGAACCGGTGTGCGAAGTAGGTGC |
| RBC | A06 | GTCTCGTGGGCTCGGAGATGTGTATAAGAGACAGATATGCCCGGTGTGCGAAGTAGGTGC |
| RBC | A07 | GTCTCGTGGGCTCGGAGATGTGTATAAGAGACAGGGTCTGACGGTGTGCGAAGTAGGTGC |
| RBC | A08 | GTCTCGTGGGCTCGGAGATGTGTATAAGAGACAGGTGAGATCGGTGTGCGAAGTAGGTGC |
| RBC | A09 | GTCTCGTGGGCTCGGAGATGTGTATAAGAGACAGTTGGCAGCGGTGTGCGAAGTAGGTGC |
| RBC | A10 | GTCTCGTGGGCTCGGAGATGTGTATAAGAGACAGATGCCTGCGGTGTGCGAAGTAGGTGC |
| RBC | A11 | GTCTCGTGGGCTCGGAGATGTGTATAAGAGACAGTCCGAAGCGGTGTGCGAAGTAGGTGC |
| RBC | A12 | GTCTCGTGGGCTCGGAGATGTGTATAAGAGACAGGGCTTACCGGTGTGCGAAGTAGGTGC |
| RBC | B01 | GTCTCGTGGGCTCGGAGATGTGTATAAGAGACAGAGTTGGCCGGTGTGCGAAGTAGGTGC |
| RBC | B02 | GTCTCGTGGGCTCGGAGATGTGTATAAGAGACAGAACGATGCGGTGTGCGAAGTAGGTGC |
| RBC | B03 | GTCTCGTGGGCTCGGAGATGTGTATAAGAGACAGACTACCGCGGTGTGCGAAGTAGGTGC |
| RBC | B04 | GTCTCGTGGGCTCGGAGATGTGTATAAGAGACAGGGTGTCTCGGTGTGCGAAGTAGGTGC |
| RBC | B05 | GTCTCGTGGGCTCGGAGATGTGTATAAGAGACAGCCAGCTTCGGTGTGCGAAGTAGGTGC |
| RBC | B06 | GTCTCGTGGGCTCGGAGATGTGTATAAGAGACAGTTAGACGCGGTGTGCGAAGTAGGTGC |
| RBC | B07 | GTCTCGTGGGCTCGGAGATGTGTATAAGAGACAGACCATACCGGTGTGCGAAGTAGGTGC |
| RBC | B08 | GTCTCGTGGGCTCGGAGATGTGTATAAGAGACAGGACGACTCGGTGTGCGAAGTAGGTGC |
| RBC | B09 | GTCTCGTGGGCTCGGAGATGTGTATAAGAGACAGGTCACCTCGGTGTGCGAAGTAGGTGC |
| RBC | B10 | GTCTCGTGGGCTCGGAGATGTGTATAAGAGACAGCGTGATGCGGTGTGCGAAGTAGGTGC |
| RBC | B11 | GTCTCGTGGGCTCGGAGATGTGTATAAGAGACAGGCTTCCTCGGTGTGCGAAGTAGGTGC |
| RBC | B12 | GTCTCGTGGGCTCGGAGATGTGTATAAGAGACAGTAGACGTCGGTGTGCGAAGTAGGTGC |
| RBC | C01 | GTCTCGTGGGCTCGGAGATGTGTATAAGAGACAGCGGACTTCGGTGTGCGAAGTAGGTGC |
| RBC | C02 | GTCTCGTGGGCTCGGAGATGTGTATAAGAGACAGACCGGAACGGTGTGCGAAGTAGGTGC |
| RBC | C03 | GTCTCGTGGGCTCGGAGATGTGTATAAGAGACAGCCGAAGTCGGTGTGCGAAGTAGGTGC |
| RBC | C04 | GTCTCGTGGGCTCGGAGATGTGTATAAGAGACAGTCACGCACGGTGTGCGAAGTAGGTGC |
| RBC | C05 | GTCTCGTGGGCTCGGAGATGTGTATAAGAGACAGATCCTCGCGGTGTGCGAAGTAGGTGC |
| RBC | C06 | GTCTCGTGGGCTCGGAGATGTGTATAAGAGACAGCGAATAGCGGTGTGCGAAGTAGGTGC |
| RBC | C07 | GTCTCGTGGGCTCGGAGATGTGTATAAGAGACAGTATCCGGCGGTGTGCGAAGTAGGTGC |
| RBC | C08 | GTCTCGTGGGCTCGGAGATGTGTATAAGAGACAGAGCAAGACGGTGTGCGAAGTAGGTGC |
| RBC | C09 | GTCTCGTGGGCTCGGAGATGTGTATAAGAGACAGTGTCGACCGGTGTGCGAAGTAGGTGC |
| RBC | C10 | GTCTCGTGGGCTCGGAGATGTGTATAAGAGACAGTTCCATGCGGTGTGCGAAGTAGGTGC |
| RBC | C11 | GTCTCGTGGGCTCGGAGATGTGTATAAGAGACAGGCAATCGCGGTGTGCGAAGTAGGTGC |
| RBC | C12 | GTCTCGTGGGCTCGGAGATGTGTATAAGAGACAGTGAGTGGCGGTGTGCGAAGTAGGTGC |
| RBC | D01 | GTCTCGTGGGCTCGGAGATGTGTATAAGAGACAGTAGGAGACGGTGTGCGAAGTAGGTGC |
| RBC | D02 | GTCTCGTGGGCTCGGAGATGTGTATAAGAGACAGAGTCAGTCGGTGTGCGAAGTAGGTGC |
| RBC | D03 | GTCTCGTGGGCTCGGAGATGTGTATAAGAGACAGGTGCTGTCGGTGTGCGAAGTAGGTGC |
| RBC | D04 | GTCTCGTGGGCTCGGAGATGTGTATAAGAGACAGCAACAACCGGTGTGCGAAGTAGGTGC |
| RBC | D05 | GTCTCGTGGGCTCGGAGATGTGTATAAGAGACAGAATAGCCCGGTGTGCGAAGTAGGTGC |
| RBC | D06 | GTCTCGTGGGCTCGGAGATGTGTATAAGAGACAGTCTGTGACGGTGTGCGAAGTAGGTGC |
| RBC | D07 | GTCTCGTGGGCTCGGAGATGTGTATAAGAGACAGTGTGGTACGGTGTGCGAAGTAGGTGC |
| RBC | D08 | GTCTCGTGGGCTCGGAGATGTGTATAAGAGACAGGCGTATGCGGTGTGCGAAGTAGGTGC |

| | | |
|---|---|---|
| RBC | D09 | GTCTCGTGGGCTCGGAGATGTGTATAAGAGACAGAGTTACGCGGTGTGCGAAGTAGGTGC |
| RBC | D10 | GTCTCGTGGGCTCGGAGATGTGTATAAGAGACAGTTCCTGCCGGTGTGCGAAGTAGGTGC |
| RBC | D11 | GTCTCGTGGGCTCGGAGATGTGTATAAGAGACAGTATGTCGCGGTGTGCGAAGTAGGTGC |
| RBC | D12 | GTCTCGTGGGCTCGGAGATGTGTATAAGAGACAGGGAGAGACGGTGTGCGAAGTAGGTGC |
| RBC | E01 | GTCTCGTGGGCTCGGAGATGTGTATAAGAGACAGCCTTAGGCGGTGTGCGAAGTAGGTGC |
| RBC | E02 | GTCTCGTGGGCTCGGAGATGTGTATAAGAGACAGTGTATCCCGGTGTGCGAAGTAGGTGC |
| RBC | E03 | GTCTCGTGGGCTCGGAGATGTGTATAAGAGACAGCAACCTGCGGTGTGCGAAGTAGGTGC |
| RBC | E04 | GTCTCGTGGGCTCGGAGATGTGTATAAGAGACAGCTGATGACGGTGTGCGAAGTAGGTGC |
| RBC | E05 | GTCTCGTGGGCTCGGAGATGTGTATAAGAGACAGAAGACAGCGGTGTGCGAAGTAGGTGC |
| RBC | E06 | GTCTCGTGGGCTCGGAGATGTGTATAAGAGACAGAGCTCGTCGGTGTGCGAAGTAGGTGC |
| RBC | E07 | GTCTCGTGGGCTCGGAGATGTGTATAAGAGACAGGATTGCGCGGTGTGCGAAGTAGGTGC |
| RBC | E08 | GTCTCGTGGGCTCGGAGATGTGTATAAGAGACAGTCCTTCACGGTGTGCGAAGTAGGTGC |
| RBC | E09 | GTCTCGTGGGCTCGGAGATGTGTATAAGAGACAGTCACAGGCGGTGTGCGAAGTAGGTGC |
| RBC | E10 | GTCTCGTGGGCTCGGAGATGTGTATAAGAGACAGAGAGCTGCGGTGTGCGAAGTAGGTGC |
| RBC | E11 | GTCTCGTGGGCTCGGAGATGTGTATAAGAGACAGCCTCTGTCGGTGTGCGAAGTAGGTGC |
| RBC | E12 | GTCTCGTGGGCTCGGAGATGTGTATAAGAGACAGCCTCGAACGGTGTGCGAAGTAGGTGC |
| RBC | F01 | GTCTCGTGGGCTCGGAGATGTGTATAAGAGACAGGTGTCTCCGGTGTGCGAAGTAGGTGC |
| RBC | F02 | GTCTCGTGGGCTCGGAGATGTGTATAAGAGACAGATTGAGGCGGTGTGCGAAGTAGGTGC |
| RBC | F03 | GTCTCGTGGGCTCGGAGATGTGTATAAGAGACAGGACAATCCGGTGTGCGAAGTAGGTGC |
| RBC | F04 | GTCTCGTGGGCTCGGAGATGTGTATAAGAGACAGCACTTGCCGGTGTGCGAAGTAGGTGC |
| RBC | F05 | GTCTCGTGGGCTCGGAGATGTGTATAAGAGACAGTGAACGCCGGTGTGCGAAGTAGGTGC |
| RBC | F06 | GTCTCGTGGGCTCGGAGATGTGTATAAGAGACAGCGTAGCACGGTGTGCGAAGTAGGTGC |
| RBC | F07 | GTCTCGTGGGCTCGGAGATGTGTATAAGAGACAGAGGTTCCCGGTGTGCGAAGTAGGTGC |
| RBC | F08 | GTCTCGTGGGCTCGGAGATGTGTATAAGAGACAGGTACACACGGTGTGCGAAGTAGGTGC |
| RBC | F09 | GTCTCGTGGGCTCGGAGATGTGTATAAGAGACAGGATAGGTCGGTGTGCGAAGTAGGTGC |
| RBC | F10 | GTCTCGTGGGCTCGGAGATGTGTATAAGAGACAGTAGCCTCCGGTGTGCGAAGTAGGTGC |
| RBC | F11 | GTCTCGTGGGCTCGGAGATGTGTATAAGAGACAGTTCAGCCCGGTGTGCGAAGTAGGTGC |
| RBC | F12 | GTCTCGTGGGCTCGGAGATGTGTATAAGAGACAGGGATTCACGGTGTGCGAAGTAGGTGC |
| RBC | G01 | GTCTCGTGGGCTCGGAGATGTGTATAAGAGACAGTGAGCCTCGGTGTGCGAAGTAGGTGC |
| RBC | G02 | GTCTCGTGGGCTCGGAGATGTGTATAAGAGACAGAACGCGACGGTGTGCGAAGTAGGTGC |
| RBC | G03 | GTCTCGTGGGCTCGGAGATGTGTATAAGAGACAGTCATTGCCGGTGTGCGAAGTAGGTGC |
| RBC | G04 | GTCTCGTGGGCTCGGAGATGTGTATAAGAGACAGAGCATCTCGGTGTGCGAAGTAGGTGC |
| RBC | G05 | GTCTCGTGGGCTCGGAGATGTGTATAAGAGACAGTTGGTCTCGGTGTGCGAAGTAGGTGC |
| RBC | G06 | GTCTCGTGGGCTCGGAGATGTGTATAAGAGACAGCAAGGATCGGTGTGCGAAGTAGGTGC |
| RBC | G07 | GTCTCGTGGGCTCGGAGATGTGTATAAGAGACAGAGACGTCCGGTGTGCGAAGTAGGTGC |
| RBC | G08 | GTCTCGTGGGCTCGGAGATGTGTATAAGAGACAGAGGTCAACGGTGTGCGAAGTAGGTGC |
| RBC | G09 | GTCTCGTGGGCTCGGAGATGTGTATAAGAGACAGATGCTACCGGTGTGCGAAGTAGGTGC |
| RBC | G10 | GTCTCGTGGGCTCGGAGATGTGTATAAGAGACAGCTCTGATCGGTGTGCGAAGTAGGTGC |
| RBC | G11 | GTCTCGTGGGCTCGGAGATGTGTATAAGAGACAGTCAAGTCCGGTGTGCGAAGTAGGTGC |
| RBC | G12 | GTCTCGTGGGCTCGGAGATGTGTATAAGAGACAGTCGAGCTCGGTGTGCGAAGTAGGTGC |
| RBC | H01 | GTCTCGTGGGCTCGGAGATGTGTATAAGAGACAGACAGTCTCGGTGTGCGAAGTAGGTGC |
| RBC | H02 | GTCTCGTGGGCTCGGAGATGTGTATAAGAGACAGCAGATACCGGTGTGCGAAGTAGGTGC |
| RBC | H03 | GTCTCGTGGGCTCGGAGATGTGTATAAGAGACAGTACGTTCCGGTGTGCGAAGTAGGTGC |

| | | |
|---|---|---|
| RBC | H04 | GTCTCGTGGGCTCGGAGATGTGTATAAGAGACAGACGGTTCCGGTGTGCGAAGTAGGTGC |
| RBC | H05 | GTCTCGTGGGCTCGGAGATGTGTATAAGAGACAGCATCGTCCGGTGTGCGAAGTAGGTGC |
| RBC | H06 | GTCTCGTGGGCTCGGAGATGTGTATAAGAGACAGTACGCATCGGTGTGCGAAGTAGGTGC |
| RBC | H07 | GTCTCGTGGGCTCGGAGATGTGTATAAGAGACAGCTTAGACCGGTGTGCGAAGTAGGTGC |
| RBC | H08 | GTCTCGTGGGCTCGGAGATGTGTATAAGAGACAGAACTGACCGGTGTGCGAAGTAGGTGC |
| RBC | H09 | GTCTCGTGGGCTCGGAGATGTGTATAAGAGACAGACTTGCACGGTGTGCGAAGTAGGTGC |
| RBC | H10 | GTCTCGTGGGCTCGGAGATGTGTATAAGAGACAGACGCGATCGGTGTGCGAAGTAGGTGC |
| RBC | H11 | GTCTCGTGGGCTCGGAGATGTGTATAAGAGACAGTCGACACCGGTGTGCGAAGTAGGTGC |
| RBC | H12 | GTCTCGTGGGCTCGGAGATGTGTATAAGAGACAGACTCAACCGGTGTGCGAAGTAGGTGC |

#### Dual-Indexing Platemaps

This section contains all platemaps for the dual indexing plates (DI plates) used in this study. The tables that follow show how the primers from the forward and reverse barcode plates (Table S2) were arrayed to produce the barcode plates. Each entry in the below platemaps follows the format "Well-Barcode Plate", where the "-" delimits the plate and well. An "F" after the delimiter indicates that the well preceding the delimiter was from the forward barcode plate ("FBC" in Table S2) and an "R" indicates that the well was from the reverse barcode plate ("RBC"). A detailed protocol for how the dual index plates were produced is given in *Preparation of evSeq Barcode Primer Mixes*, above.

**Table S3.** Platemap for DI01 used in this study.

| **DI01** | 01 | 02 | 03 | 04 | 05 | 06 | 07 | 08 | 09 | 10 | 11 | 12 |
|---|---|---|---|---|---|---|---|---|---|---|---|---|
| A | A01-F, A01-R | A02-F, A02-R | A03-F, A03-R | A04-F, A04-R | A05-F, A05-R | A06-F, A06-R | A07-F, A07-R | A08-F, A08-R | A09-F, A09-R | A10-F, A10-R | A11-F, A11-R | A12-F, A12-R |
| B | B01-F, B01-R | B02-F, B02-R | B03-F, B03-R | B04-F, B04-R | B05-F, B05-R | B06-F, B06-R | B07-F, B07-R | B08-F, B08-R | B09-F, B09-R | B10-F, B10-R | B11-F, B11-R | B12-F, B12-R |
| C | C01-F, C01-R | C02-F, C02-R | C03-F, C03-R | C04-F, C04-R | C05-F, C05-R | C06-F, C06-R | C07-F, C07-R | C08-F, C08-R | C09-F, C09-R | C10-F, C10-R | C11-F, C11-R | C12-F, C12-R |
| D | D01-F, D01-R | D02-F, D02-R | D03-F, D03-R | D04-F, D04-R | D05-F, D05-R | D06-F, D06-R | D07-F, D07-R | D08-F, D08-R | D09-F, D09-R | D10-F, D10-R | D11-F, D11-R | D12-F, D12-R |
| E | E01-F, E01-R | E02-F, E02-R | E03-F, E03-R | E04-F, E04-R | E05-F, E05-R | E06-F, E06-R | E07-F, E07-R | E08-F, E08-R | E09-F, E09-R | E10-F, E10-R | E11-F, E11-R | E12-F, E12-R |
| F | F01-F, F01-R | F02-F, F02-R | F03-F, F03-R | F04-F, F04-R | F05-F, F05-R | F06-F, F06-R | F07-F, F07-R | F08-F, F08-R | F09-F, F09-R | F10-F, F10-R | F11-F, F11-R | F12-F, F12-R |
| G | G01-F, G01-R | G02-F, G02-R | G03-F, G03-R | G04-F, G04-R | G05-F, G05-R | G06-F, G06-R | G07-F, G07-R | G08-F, G08-R | G09-F, G09-R | G10-F, G10-R | G11-F, G11-R | G12-F, G12-R |
| H | H01-F, H01-R | H02-F, H02-R | H03-F, H03-R | H04-F, H04-R | H05-F, H05-R | H06-F, H06-R | H07-F, H07-R | H08-F, H08-R | H09-F, H09-R | H10-F, H10-R | H11-F, H11-R | H12-F, H12-R |

**Table S4.** Platemap for DI02 used in this study.

| **DI02** | 01 | 02 | 03 | 04 | 05 | 06 | 07 | 08 | 09 | 10 | 11 | 12 |
|---|---|---|---|---|---|---|---|---|---|---|---|---|
| A | A01-F, H01-R | A02-F, H02-R | A03-F, H03-R | A04-F, H04-R | A05-F, H05-R | A06-F, H06-R | A07-F, H07-R | A08-F, H08-R | A09-F, H09-R | A10-F, H10-R | A11-F, H11-R | A12-F, H12-R |
| B | B01-F, A01-R | B02-F, A02-R | B03-F, A03-R | B04-F, A04-R | B05-F, A05-R | B06-F, A06-R | B07-F, A07-R | B08-F, A08-R | B09-F, A09-R | B10-F, A10-R | B11-F, A11-R | B12-F, A12-R |
| C | C01-F, B01-R | C02-F, B02-R | C03-F, B03-R | C04-F, B04-R | C05-F, B05-R | C06-F, B06-R | C07-F, B07-R | C08-F, B08-R | C09-F, B09-R | C10-F, B10-R | C11-F, B11-R | C12-F, B12-R |
| D | D01-F, C01-R | D02-F, C02-R | D03-F, C03-R | D04-F, C04-R | D05-F, C05-R | D06-F, C06-R | D07-F, C07-R | D08-F, C08-R | D09-F, C09-R | D10-F, C10-R | D11-F, C11-R | D12-F, C12-R |
| E | E01-F, D01-R | E02-F, D02-R | E03-F, D03-R | E04-F, D04-R | E05-F, D05-R | E06-F, D06-R | E07-F, D07-R | E08-F, D08-R | E09-F, D09-R | E10-F, D10-R | E11-F, D11-R | E12-F, D12-R |
| F | F01-F, E01-R | F02-F, E02-R | F03-F, E03-R | F04-F, E04-R | F05-F, E05-R | F06-F, E06-R | F07-F, E07-R | F08-F, E08-R | F09-F, E09-R | F10-F, E10-R | F11-F, E11-R | F12-F, E12-R |
| G | G01-F, F01-R | G02-F, F02-R | G03-F, F03-R | G04-F, F04-R | G05-F, F05-R | G06-F, F06-R | G07-F, F07-R | G08-F, F08-R | G09-F, F09-R | G10-F, F10-R | G11-F, F11-R | G12-F, F12-R |
| H | H01-F, G01-R | H02-F, G02-R | H03-F, G03-R | H04-F, G04-R | H05-F, G05-R | H06-F, G06-R | H07-F, G07-R | H08-F, G08-R | H09-F, G09-R | H10-F, G10-R | H11-F, G11-R | H12-F, G12-R |

**Table S5.** Platemap for DI03 used in this study.

| **DI03** | 01 | 02 | 03 | 04 | 05 | 06 | 07 | 08 | 09 | 10 | 11 | 12 |
|---|---|---|---|---|---|---|---|---|---|---|---|---|
| A | A01-F, G01-R | A02-F, G02-R | A03-F, G03-R | A04-F, G04-R | A05-F, G05-R | A06-F, G06-R | A07-F, G07-R | A08-F, G08-R | A09-F, G09-R | A10-F, G10-R | A11-F, G11-R | A12-F, G12-R |
| B | B01-F, H01-R | B02-F, H02-R | B03-F, H03-R | B04-F, H04-R | B05-F, H05-R | B06-F, H06-R | B07-F, H07-R | B08-F, H08-R | B09-F, H09-R | B10-F, H10-R | B11-F, H11-R | B12-F, H12-R |
| C | C01-F, A01-R | C02-F, A02-R | C03-F, A03-R | C04-F, A04-R | C05-F, A05-R | C06-F, A06-R | C07-F, A07-R | C08-F, A08-R | C09-F, A09-R | C10-F, A10-R | C11-F, A11-R | C12-F, A12-R |
| D | D01-F, B01-R | D02-F, B02-R | D03-F, B03-R | D04-F, B04-R | D05-F, B05-R | D06-F, B06-R | D07-F, B07-R | D08-F, B08-R | D09-F, B09-R | D10-F, B10-R | D11-F, B11-R | D12-F, B12-R |
| E | E01-F, C01-R | E02-F, C02-R | E03-F, C03-R | E04-F, C04-R | E05-F, C05-R | E06-F, C06-R | E07-F, C07-R | E08-F, C08-R | E09-F, C09-R | E10-F, C10-R | E11-F, C11-R | E12-F, C12-R |
| F | F01-F, D01-R | F02-F, D02-R | F03-F, D03-R | F04-F, D04-R | F05-F, D05-R | F06-F, D06-R | F07-F, D07-R | F08-F, D08-R | F09-F, D09-R | F10-F, D10-R | F11-F, D11-R | F12-F, D12-R |
| G | G01-F, E01-R | G02-F, E02-R | G03-F, E03-R | G04-F, E04-R | G05-F, E05-R | G06-F, E06-R | G07-F, E07-R | G08-F, E08-R | G09-F, E09-R | G10-F, E10-R | G11-F, E11-R | G12-F, E12-R |
| H | H01-F, F01-R | H02-F, F02-R | H03-F, F03-R | H04-F, F04-R | H05-F, F05-R | H06-F, F06-R | H07-F, F07-R | H08-F, F08-R | H09-F, F09-R | H10-F, F10-R | H11-F, F11-R | H12-F, F12-R |

**Table S6.** Platemap for DI04 used in this study.

| **DI04** | 01 | 02 | 03 | 04 | 05 | 06 | 07 | 08 | 09 | 10 | 11 | 12 |
|---|---|---|---|---|---|---|---|---|---|---|---|---|
| A | A01-F, F01-R | A02-F, F02-R | A03-F, F03-R | A04-F, F04-R | A05-F, F05-R | A06-F, F06-R | A07-F, F07-R | A08-F, F08-R | A09-F, F09-R | A10-F, F10-R | A11-F, F11-R | A12-F, F12-R |
| B | B01-F, G01-R | B02-F, G02-R | B03-F, G03-R | B04-F, G04-R | B05-F, G05-R | B06-F, G06-R | B07-F, G07-R | B08-F, G08-R | B09-F, G09-R | B10-F, G10-R | B11-F, G11-R | B12-F, G12-R |
| C | C01-F, H01-R | C02-F, H02-R | C03-F, H03-R | C04-F, H04-R | C05-F, H05-R | C06-F, H06-R | C07-F, H07-R | C08-F, H08-R | C09-F, H09-R | C10-F, H10-R | C11-F, H11-R | C12-F, H12-R |
| D | D01-F, A01-R | D02-F, A02-R | D03-F, A03-R | D04-F, A04-R | D05-F, A05-R | D06-F, A06-R | D07-F, A07-R | D08-F, A08-R | D09-F, A09-R | D10-F, A10-R | D11-F, A11-R | D12-F, A12-R |
| E | E01-F, B01-R | E02-F, B02-R | E03-F, B03-R | E04-F, B04-R | E05-F, B05-R | E06-F, B06-R | E07-F, B07-R | E08-F, B08-R | E09-F, B09-R | E10-F, B10-R | E11-F, B11-R | E12-F, B12-R |
| F | F01-F, C01-R | F02-F, C02-R | F03-F, C03-R | F04-F, C04-R | F05-F, C05-R | F06-F, C06-R | F07-F, C07-R | F08-F, C08-R | F09-F, C09-R | F10-F, C10-R | F11-F, C11-R | F12-F, C12-R |
| G | G01-F, D01-R | G02-F, D02-R | G03-F, D03-R | G04-F, D04-R | G05-F, D05-R | G06-F, D06-R | G07-F, D07-R | G08-F, D08-R | G09-F, D09-R | G10-F, D10-R | G11-F, D11-R | G12-F, D12-R |
| H | H01-F, E01-R | H02-F, E02-R | H03-F, E03-R | H04-F, E04-R | H05-F, E05-R | H06-F, E06-R | H07-F, E07-R | H08-F, E08-R | H09-F, E09-R | H10-F, E10-R | H11-F, E11-R | H12-F, E12-R |

**Table S7.** Platemap for DI05 used in this study.

| **DI05** | 01 | 02 | 03 | 04 | 05 | 06 | 07 | 08 | 09 | 10 | 11 | 12 |
|---|---|---|---|---|---|---|---|---|---|---|---|---|
| A | A01-F, E01-R | A02-F, E02-R | A03-F, E03-R | A04-F, E04-R | A05-F, E05-R | A06-F, E06-R | A07-F, E07-R | A08-F, E08-R | A09-F, E09-R | A10-F, E10-R | A11-F, E11-R | A12-F, E12-R |
| B | B01-F, F01-R | B02-F, F02-R | B03-F, F03-R | B04-F, F04-R | B05-F, F05-R | B06-F, F06-R | B07-F, F07-R | B08-F, F08-R | B09-F, F09-R | B10-F, F10-R | B11-F, F11-R | B12-F, F12-R |
| C | C01-F, G01-R | C02-F, G02-R | C03-F, G03-R | C04-F, G04-R | C05-F, G05-R | C06-F, G06-R | C07-F, G07-R | C08-F, G08-R | C09-F, G09-R | C10-F, G10-R | C11-F, G11-R | C12-F, G12-R |
| D | D01-F, H01-R | D02-F, H02-R | D03-F, H03-R | D04-F, H04-R | D05-F, H05-R | D06-F, H06-R | D07-F, H07-R | D08-F, H08-R | D09-F, H09-R | D10-F, H10-R | D11-F, H11-R | D12-F, H12-R |
| E | E01-F, A01-R | E02-F, A02-R | E03-F, A03-R | E04-F, A04-R | E05-F, A05-R | E06-F, A06-R | E07-F, A07-R | E08-F, A08-R | E09-F, A09-R | E10-F, A10-R | E11-F, A11-R | E12-F, A12-R |
| F | F01-F, B01-R | F02-F, B02-R | F03-F, B03-R | F04-F, B04-R | F05-F, B05-R | F06-F, B06-R | F07-F, B07-R | F08-F, B08-R | F09-F, B09-R | F10-F, B10-R | F11-F, B11-R | F12-F, B12-R |
| G | G01-F, C01-R | G02-F, C02-R | G03-F, C03-R | G04-F, C04-R | G05-F, C05-R | G06-F, C06-R | G07-F, C07-R | G08-F, C08-R | G09-F, C09-R | G10-F, C10-R | G11-F, C11-R | G12-F, C12-R |
| H | H01-F, D01-R | H02-F, D02-R | H03-F, D03-R | H04-F, D04-R | H05-F, D05-R | H06-F, D06-R | H07-F, D07-R | H08-F, D08-R | H09-F, D09-R | H10-F, D10-R | H11-F, D11-R | H12-F, D12-R |

**Table S8.** Platemap for DI06 used in this study.

| **DI06** | 01 | 02 | 03 | 04 | 05 | 06 | 07 | 08 | 09 | 10 | 11 | 12 |
|---|---|---|---|---|---|---|---|---|---|---|---|---|
| A | A01-F, D01-R | A02-F, D02-R | A03-F, D03-R | A04-F, D04-R | A05-F, D05-R | A06-F, D06-R | A07-F, D07-R | A08-F, D08-R | A09-F, D09-R | A10-F, D10-R | A11-F, D11-R | A12-F, D12-R |
| B | B01-F, E01-R | B02-F, E02-R | B03-F, E03-R | B04-F, E04-R | B05-F, E05-R | B06-F, E06-R | B07-F, E07-R | B08-F, E08-R | B09-F, E09-R | B10-F, E10-R | B11-F, E11-R | B12-F, E12-R |
| C | C01-F, F01-R | C02-F, F02-R | C03-F, F03-R | C04-F, F04-R | C05-F, F05-R | C06-F, F06-R | C07-F, F07-R | C08-F, F08-R | C09-F, F09-R | C10-F, F10-R | C11-F, F11-R | C12-F, F12-R |
| D | D01-F, G01-R | D02-F, G02-R | D03-F, G03-R | D04-F, G04-R | D05-F, G05-R | D06-F, G06-R | D07-F, G07-R | D08-F, G08-R | D09-F, G09-R | D10-F, G10-R | D11-F, G11-R | D12-F, G12-R |
| E | E01-F, H01-R | E02-F, H02-R | E03-F, H03-R | E04-F, H04-R | E05-F, H05-R | E06-F, H06-R | E07-F, H07-R | E08-F, H08-R | E09-F, H09-R | E10-F, H10-R | E11-F, H11-R | E12-F, H12-R |
| F | F01-F, A01-R | F02-F, A02-R | F03-F, A03-R | F04-F, A04-R | F05-F, A05-R | F06-F, A06-R | F07-F, A07-R | F08-F, A08-R | F09-F, A09-R | F10-F, A10-R | F11-F, A11-R | F12-F, A12-R |
| G | G01-F, B01-R | G02-F, B02-R | G03-F, B03-R | G04-F, B04-R | G05-F, B05-R | G06-F, B06-R | G07-F, B07-R | G08-F, B08-R | G09-F, B09-R | G10-F, B10-R | G11-F, B11-R | G12-F, B12-R |
| H | H01-F, C01-R | H02-F, C02-R | H03-F, C03-R | H04-F, C04-R | H05-F, C05-R | H06-F, C06-R | H07-F, C07-R | H08-F, C08-R | H09-F, C09-R | H10-F, C10-R | H11-F, C11-R | H12-F, C12-R |

**Table S9.** Platemap for DI07 used in this study.

| **DI07** | 01 | 02 | 03 | 04 | 05 | 06 | 07 | 08 | 09 | 10 | 11 | 12 |
|---|---|---|---|---|---|---|---|---|---|---|---|---|
| A | A01-F, C01-R | A02-F, C02-R | A03-F, C03-R | A04-F, C04-R | A05-F, C05-R | A06-F, C06-R | A07-F, C07-R | A08-F, C08-R | A09-F, C09-R | A10-F, C10-R | A11-F, C11-R | A12-F, C12-R |
| B | B01-F, D01-R | B02-F, D02-R | B03-F, D03-R | B04-F, D04-R | B05-F, D05-R | B06-F, D06-R | B07-F, D07-R | B08-F, D08-R | B09-F, D09-R | B10-F, D10-R | B11-F, D11-R | B12-F, D12-R |
| C | C01-F, E01-R | C02-F, E02-R | C03-F, E03-R | C04-F, E04-R | C05-F, E05-R | C06-F, E06-R | C07-F, E07-R | C08-F, E08-R | C09-F, E09-R | C10-F, E10-R | C11-F, E11-R | C12-F, E12-R |
| D | D01-F, F01-R | D02-F, F02-R | D03-F, F03-R | D04-F, F04-R | D05-F, F05-R | D06-F, F06-R | D07-F, F07-R | D08-F, F08-R | D09-F, F09-R | D10-F, F10-R | D11-F, F11-R | D12-F, F12-R |
| E | E01-F, G01-R | E02-F, G02-R | E03-F, G03-R | E04-F, G04-R | E05-F, G05-R | E06-F, G06-R | E07-F, G07-R | E08-F, G08-R | E09-F, G09-R | E10-F, G10-R | E11-F, G11-R | E12-F, G12-R |
| F | F01-F, H01-R | F02-F, H02-R | F03-F, H03-R | F04-F, H04-R | F05-F, H05-R | F06-F, H06-R | F07-F, H07-R | F08-F, H08-R | F09-F, H09-R | F10-F, H10-R | F11-F, H11-R | F12-F, H12-R |
| G | G01-F, A01-R | G02-F, A02-R | G03-F, A03-R | G04-F, A04-R | G05-F, A05-R | G06-F, A06-R | G07-F, A07-R | G08-F, A08-R | G09-F, A09-R | G10-F, A10-R | G11-F, A11-R | G12-F, A12-R |
| H | H01-F, B01-R | H02-F, B02-R | H03-F, B03-R | H04-F, B04-R | H05-F, B05-R | H06-F, B06-R | H07-F, B07-R | H08-F, B08-R | H09-F, B09-R | H10-F, B10-R | H11-F, B11-R | H12-F, B12-R |

**Table S10.** Platemap for DI08 used in this study.

| **DI08** | 01 | 02 | 03 | 04 | 05 | 06 | 07 | 08 | 09 | 10 | 11 | 12 |
|---|---|---|---|---|---|---|---|---|---|---|---|---|
| A | A01-F, B01-R | A02-F, B02-R | A03-F, B03-R | A04-F, B04-R | A05-F, B05-R | A06-F, B06-R | A07-F, B07-R | A08-F, B08-R | A09-F, B09-R | A10-F, B10-R | A11-F, B11-R | A12-F, B12-R |
| B | B01-F, C01-R | B02-F, C02-R | B03-F, C03-R | B04-F, C04-R | B05-F, C05-R | B06-F, C06-R | B07-F, C07-R | B08-F, C08-R | B09-F, C09-R | B10-F, C10-R | B11-F, C11-R | B12-F, C12-R |
| C | C01-F, D01-R | C02-F, D02-R | C03-F, D03-R | C04-F, D04-R | C05-F, D05-R | C06-F, D06-R | C07-F, D07-R | C08-F, D08-R | C09-F, D09-R | C10-F, D10-R | C11-F, D11-R | C12-F, D12-R |
| D | D01-F, E01-R | D02-F, E02-R | D03-F, E03-R | D04-F, E04-R | D05-F, E05-R | D06-F, E06-R | D07-F, E07-R | D08-F, E08-R | D09-F, E09-R | D10-F, E10-R | D11-F, E11-R | D12-F, E12-R |
| E | E01-F, F01-R | E02-F, F02-R | E03-F, F03-R | E04-F, F04-R | E05-F, F05-R | E06-F, F06-R | E07-F, F07-R | E08-F, F08-R | E09-F, F09-R | E10-F, F10-R | E11-F, F11-R | E12-F, F12-R |
| F | F01-F, G01-R | F02-F, G02-R | F03-F, G03-R | F04-F, G04-R | F05-F, G05-R | F06-F, G06-R | F07-F, G07-R | F08-F, G08-R | F09-F, G09-R | F10-F, G10-R | F11-F, G11-R | F12-F, G12-R |
| G | G01-F, H01-R | G02-F, H02-R | G03-F, H03-R | G04-F, H04-R | G05-F, H05-R | G06-F, H06-R | G07-F, H07-R | G08-F, H08-R | G09-F, H09-R | G10-F, H10-R | G11-F, H11-R | G12-F, H12-R |
| H | H01-F, A01-R | H02-F, A02-R | H03-F, A03-R | H04-F, A04-R | H05-F, A05-R | H06-F, A06-R | H07-F, A07-R | H08-F, A08-R | H09-F, A09-R | H10-F, A10-R | H11-F, A11-R | H12-F, A12-R |

#### Supplemental Tables

**Table S11.** evSeq captures off-target mutations. This table is derived from the “AminoAcids_Coupled_Max.csv” output file from evSeq for the TrpB run, and shows all confident (defined as >0.80 alignment frequency and >10 total reads) unexpected mutations captured by evSeq; some columns have been removed. Note in the “VariantCombo” column that the amino acid at the expected mutagenized position has a “?” as the original amino acid—this is because the evSeq run generating this data was told the variable positions with the “NNN” convention. For unexpected variable positions, both the original amino acid and the new amino acid are shown.

| IndexPlate | Plate | Well | VariantCombo | AlignmentFrequency | WellSeqDepth |
|---|---|---|---|---:|---:|
| DI02 | Lib2_118X | E03 | ?118V_D164G | 0.964286 | 28 |
| DI04 | Lib4_166X | B02 | P154S_?166Q | 0.977011 | 87 |
| DI08 | Lib8_301X | H11 | G250D_?301L | 0.99537 | 216 |

#### Supplemental References

(1) Kille, S.; Acevedo-Rocha, C. G.; Parra, L. P.; Zhang, Z.-G.; Opperman, D. J.; Reetz, M. T.; Acevedo, J. P. Reducing Codon Redundancy and Screening Effort of Combinatorial Protein Libraries Created by Saturation Mutagenesis. *ACS Synth. Biol.* **2013**, *2* (2), 83–92.

(2) Gibson, D. G.; Young, L.; Chuang, R. Y.; Venter, J. C.; Hutchison, C. A.; Smith, H. O. Enzymatic Assembly of DNA Molecules up to Several Hundred Kilobases. *Nat. Methods* **2009**, *6* (5), 343–345.

(3) Rix, G.; Watkins-Dulaney, E. J.; Almhjell, P. J.; Boville, C. E.; Arnold, F. H.; Liu, C. C. Scalable Continuous Evolution for the Generation of Diverse Enzyme Variants Encompassing Promiscuous Activities. *Nat. Commun.* **2020**, *11* (1), 1–11.
